# Diverse DNA modification in marine prokaryotic and viral communities

**DOI:** 10.1101/2021.05.08.442635

**Authors:** Satoshi Hiraoka, Tomomi Sumida, Miho Hirai, Atsushi Toyoda, Shinsuke Kawagucci, Taichi Yokokawa, Takuro Nunoura

## Abstract

Chemical modifications of DNA, including methylation, play an important role in prokaryotes and viruses. However, our knowledge of the modification systems in environmental microbial communities, typically dominated by members not yet cultured, is limited. Here, we conducted ‘metaepigenomic’ analyses by single-molecule real-time sequencing of marine microbial communities. In total, 233 and 163 metagenomic assembly genomes (MAGs) were constructed from diverse prokaryotes and viruses, respectively, and 220 modified motifs and 276 DNA methyltransferases (MTases) were identified. Most of the MTases were not associated with the defense mechanism. The MTase-motif correspondence found in the MAGs revealed 10 novel pairs, and experimentally confirmed the catalytic specificities of the MTases. We revealed novel alternative motifs in the methylation system that are highly conserved in Alphaproteobacteria, illuminating the co-evolutionary history of the methylation system and host genome. Our findings highlight diverse unexplored DNA modifications that potentially affect the ecology and evolution of prokaryotes and viruses.

## Introduction

DNA chemical modifications are found in diverse prokaryotes and viruses as well as eukaryotes. DNA methylation, a representative DNA modification, is catalyzed by DNA methyltransferase (MTase), whereas S-adenosylmethionine (SAM) provides a methyl group (*1*). In prokaryotes, three types of methylation (i.e., N6-methyladenine (m6A), 5-methylcytosine (m5C), and N4-methylcytosine (m4C)) have been investigated in detail (*2*). DNA methylation plays a role in the regulation of gene expression and mismatch DNA repair (*3–5*). Subsequently, the systems function on physiological machineries, including asymmetric cell division (*6, 7*), ultraviolet (UV) tolerance (*8*), motility (*9*), and virulence of pathogens (*10–12*). DNA methylation also facilitates cell protection from invasion of extracellular DNA, such as viruses and plasmids, known as restriction-modification (RM) systems (*13*). To overcome the defense system, some viruses possess MTases and modify their genomic DNA to escape the host RM system (*2*). Moreover, frequent gene duplication, loss, and horizontal gene transfer within and between domains have been revealed during the evolution of prokaryotes (*14*). In addition to methylation, other epigenetic modifications, such as phosphorothioate, have recently been shown to have a significant effect on cells, including maintenance of cellular redox homeostasis and epigenetic regulation (*15*). Research interests in various epigenomic systems in diverse prokaryotes and viruses are increasing because of their importance in microbial physiology, genetics, evolution, and disease pathogenicity (*16–18*). However, current knowledge of prokaryotic and viral epigenomics mostly relies on studies with culturable strains, although the majority of microbes have not yet been cultured.

The recent development of single-molecule real-time (SMRT) sequencing technology, one of the feasible methods to detect DNA modification, and its implementation in PacBio sequencing platforms, has revealed an array of DNA modifications of prokaryotic (*19–24*) and viral strains (*25, 26*). The capability of long reads with few context-specific biases (*e.g*., GC bias) (*27*) allows circler consensus sequencing (CCS) method to generate highly accurate high-fidelity (HiFi) reads through error collection with multiple ‘subreads’ sequences in every single read (*28*). Based on the innovative SMRT sequencing technique, we applied culture-independent shotgun metagenomic and epigenomic analyses on freshwater microbial communities to reveal the vast DNA modification systems in nature, and established ‘metaepigenomics’ (*29*). Apart from PacBio, nanopore sequencing platforms produced by Oxford Nanopore Technologies (ONT) achieve longer reads that potentially improve metagenomic assembly with high diversity (*30*). Thus, a hybrid approach with HiFi and ONT reads is an ideal way to improve metaepigenomic analysis with accurate identification of modifications from organisms in highly diverse microbial communities.

Here, we conducted a metaepigenomic analysis of pelagic microbial communities using SMRT sequencing technology to reveal the epigenomic characteristics of diverse marine prokaryotes and viruses whose epigenomic states have not been well described. The diverse DNA modifications were successfully characterized on numerous metagenomic assembled genomes (MAGs) from both prokaryotes and viruses obtained by a combination of PacBio Sequel, ONT GridION, and Illumina Miseq sequencing platforms. Our computational prediction and experimental assay confirmed several MTases responsible for the detected methylated motifs, including novel ones. In particular, a highly conserved methylation system with varied specificity was found in Alphaproteobacteria, suggesting co-evolution between the methylation systems and their host genomes.

## Results and discussions

### Seawater sampling

Four seawater samples were collected from epipelagic (5 and 90 mbsl) and mesopelagic (200 and 300 mbsl) layers from two closely located stations in the Pacific Ocean (referred to as CM1_5m, Ct9H_90m, CM1_200m, and Ct9H_300m) (Fig. S1a, Table S1). The hydrographic properties were distinct between the epipelagic and mesopelagic zones (*31*) (Fig. S1b). Subsurface chlorophyll *a* maximum appeared at 60–100 mbsl during the samplings. The densities of prokaryotic cells and virus-like particles were highest in the uppermost layer and decreased with increasing water depth. The virus-to-prokaryote ratio increased with increasing water depth.

### Shotgun sequencing

PacBio Sequel produced 16–21 million (75–104 Gb) subreads from each sample (Table S2). The CCS analysis produced 0.66–1.1 million HiFi reads with >99% accuracy and average lengths ranging from 4311 to 4926 bp (Figs. S2a-d). The HiFi reads were estimated to cover 42–63% of the community diversity in each sample (Fig. S3). In addition to the SMRT sequencing, we conducted shotgun sequencing of CM1_5m using GridION and obtained 25 million (67 Gb) ONT reads (Table S2). The average length of the ONT reads (2734 ± 2013 bp) was comparatively shorter than the HiFi reads with high deviation, likely due to the methods used for DNA extraction based on bead-beating technique, although N50 reached 3.5 kb and the longest read achieved was 200 kb length (Fig. S2e). Illumina MiSeq reads were also obtained for each sample (Table S2).

### HiFi read analysis

Taxonomic assignment of the HiFi reads was performed using Kaiju (*32*) with the NCBI nr database (*33*) (Figs. S4a-c), and 83–89% and 3–12% of the HiFi reads were assigned to Bacteria and Archaea, respectively. Only 2–8% and 0.4–3.7% of HiFi reads were assigned to Eukaryota and Viruses, respectively. Assignment ratios of prokaryotes were 85–91%, 31–74%, and 21–53% at the phylum, order, and genus levels, respectively. Similar assignment was obtained by using the Global Ocean Reference Genomes Tropics (GORG-Tropics) database, which is composed of single-cell amplified genomes from pelagic surface layers (*34*) (Figs. S4d-f). Taxonomic compositions of full-length 16S rRNA gene sequences in the HiFi reads using BLASTN (*35*) against the SILVA database (*36*) were also consistent with the protein-based assignments using nr or GORG-Tropics (Figs. S4g-i). These similarities verify that the profile estimated using Kaiju with nr (Figs. S4a-c) is appropriate for further analysis.

Within the prokaryotes, Proteobacteria predominated in all samples at the phylum level (36–52%). Cyanobacteria was the next most abundant phylum in the epipelagic waters, CM1_5m (19%) and Ct9H_90m (27%), where photosynthesis is active. In the mesopelagic CM1_200m and Ct9H_300m communities, Chloroflexi (8.0 and 8.6%, respectively) and Thaumarchaeota (6.1 and 7.9%) were the dominant phyla. Actinobacteria (4.4–5.0%), Bacteroidetes (2.3–5.4%), and Euryarchaeota (3.1–5.3%) were abundant in all samples. The ratio of Archaea was relatively low in the epipelagic samples (3.3 and 5.1%) while high in the mesopelagic samples (11.8 and 12.2%). Within eukaryotes, the dominant phylum was Chlorophyta (0.2–3.2%), followed by Haptista (0.2–1.2%) and Ascomycota (0.5–0.7%). The highest abundance of eukaryotes was observed in Ct9H_90m from the chlorophyll *a* maximum zone. Viral reads were abundant in the epipelagic samples (4.0–4.7%) and scarce in the mesopelagic samples (0.66–0.67%). Among the viruses, Myoviridae was the most abundant family, followed by Siphoviridae, Phycodnaviridae, and Podoviridae. Myoviridae, Siphoviridae, and Podoviridae belong to the order Caudovirales, known as prokaryotic viruses (*37*). In contrast, Phycodnaviridae, a member of giant viruses with large capsids, primarily infects eukaryotic algae, including members of Chlorophyta (*38*). Because major floating viruses passed through the used 0.22-μm membrane filters, most of the viral reads were expected to be derived from lytic viruses replicating in host cells, lysogens within host genomes, or extracellular giant viruses. Viruses without double-stranded DNA (*i.e*., single-stranded DNA and RNA viruses) were not observed due to the experimental method employed. Overall, the taxonomic compositions were consistent with those of previous studies (*39–42*).

The abundance of genes related to DNA methylation and RM system in the samples were investigated by systematic annotation of MTase and restriction enzyme (REase) genes on HiFi reads using REBASE Gold Standard database (*43*). Generally, genes assigned to MTase (M), REase (R), and protein fused with the MTase and REase domains (RM) showed similar compositions among the microbial communities, and their relative abundances decreased slightly with increasing water depth (Fig. 1a). Within the MTase proteins (i.e., M and RM), Type II predominated, accounting for 76.5–78.6% in each sample (Fig. 1b). The relative abundances of Type I (11.2–12.1%) and III (3.4–5.9%) were approximately 2–3 times lower than those identified in the genomes of prokaryotic isolates, reported as 27% and 8%, respectively (*20*). Among the detected MTases, the most abundant modification type was m6A (56.9–62.9%), followed by m4C (15.6–19.6%) and m5C (14.6–19.6%) (Fig. 1c).

**Fig. 1.**
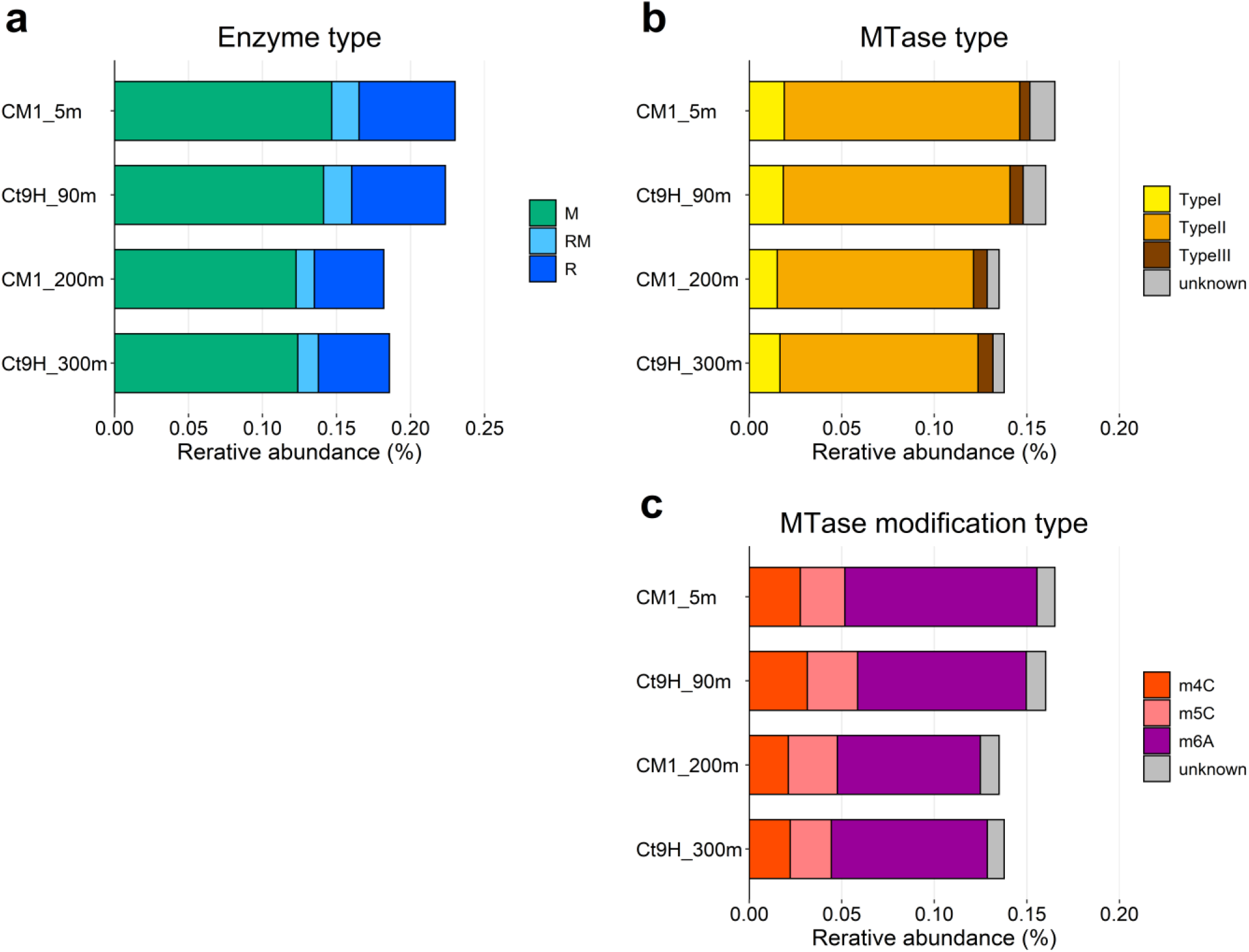
Relative abundances of DNA restriction and modification enzyme genes. CDSs predicted from HiFi reads were used in this analysis. **a** Distribution of RM types: DNA methyltransferase (MTase; M), Restriction endonuclease (REase; R), and protein fused with M and R domains (RM). **b** Distribution of MTase types. **c** Distribution of modification types.

### Metagenomic assembly and genome binning

The HiFi and ONT reads were assembled into 7829–29391 contigs (Table S3, Fig. S5). The total length of the assembled contigs from the HiFi reads (Ct9H_90m, CM1_200m, and Ct9H_300m) ranged from 83–123 Mb, the N50 values were 11–13 kb, and the length of the longest contigs was 308–546 kb. In contrast, the contigs of CM1_5m assembled from ONT reads after polishing showed outperformed statistics that reached a total length of 524 Mb with a 29 kb N50 value, likely due to the number of ONT reads longer than the HiFi reads.

The contigs were binned into a total of 233 prokaryotic MAGs (P-MAGs) (Table S3, Data S1). Among the contigs of CM1_5m, 1165 (3.96%) were assigned to 130 P-MAGs. From Ct9H_90m, CM1_200, and Ct9H_300m, 1009 (12.9%), 3366 (31.1%), and 2761 (33.9%) contigs assembled from the HiFi reads were binned into 30, 41, and 32 P-MAGs, respectively. In total, 9.4–31.5% of the HiFi reads were re-mapped to the P-MAGs for each sample. The completeness of the P-MAGs was 28.6% on average, and the highest value was 98.8%. The estimated contamination levels were low (1.2%, on average). N50 ranged from 6.4 kb to 1.42 Mb. Subread coverages ranged from 31.4–3305.7× per P-MAGs, and 218 P-MAGs (94%) showed >50×, which is sufficient to detect m6A and m4C modifications using SMRT sequencing according to the manufacture’s instruction.

A total of 233 P-MAGs spanned 14 phyla (Data S1). Among the P-MAGs, we predicted full-length 16S rRNA gene sequences from 64 P-MAGs. No assignment was obtained for 10 (4.3%) and 56 (24%) P-MAGs at the phylum and class levels, respectively, likely owing to the existence of vast phylogenetically undescribed lineages. A total of 110 (47%) P-MAGs belonged to the phylum Proteobacteria, including 41 and 52 P-MAGs assigned to the class Alphaproteobacteria and Gammaproteobacteria, respectively. Most of the Alphaproteobacteria P-MAGs were assigned to the order *Candidatus* (*Ca*.) Pelagibacterales, known as SAR11, which is one of the most abundant groups in pelagic ecosystems (*44*). P-MAGs affiliated to other dominant phyla such as Chloroflexi (18 P-MAGs); *Ca*. Marinimicrobia (18 P-MAGs), Acidobacteria (16 P-MAGs), Bacteroidetes (15 P-MAGs), Verrucomicrobia (5 P-MAGs), and Cyanobacteria (5 P-MAGs) were also identified. Among the Chloroflexi P-MAGs, six 16S rRNA gene sequences were identified, all of which were assigned to SAR202 clade. P-MAGs assigned to Archaea were composed of two phyla: Euryarchaeota (27 P-MAGs) and Thaumarchaeota (3 P-MAGs).

In addition to the prokaryotic MAGs, a total of 163 viral MAGs (V-MAGs) were retrieved from the contigs (Data S1). Most (98%) of the V-MAGs were retrieved from CM1_5m, while three, one, and zero V-MAGs were from Ct9H_90m, CM1_200m, and Ct9H_300m, respectively. This distinct difference between CM1_5m and the others likely resulted from the variance of virus abundance and the outstanding efficiencies of metagenomic assembly using ONT reads only available from CM1_5m (Figs. S4a, S5, and Table S3). The lengths of the V-MAGs ranged from 8–248 kb and 64 kb on average. Subread coverages ranged from 1.5–894.8× and 103 P-MAGs (63%) were >50×. In total, 4.72% of the HiFi reads were re-mapped to the V-MAGs in CM1_5m. Nine V-MAGs were identified as proviruses. The most abundant order was Siphoviridae (26 V-MAGs), followed by Myoviridae (14 V-MAGs), Phycodnaviridae (2 V-MAGs), and Podoviridae (2 V-MAGs), while most of the 119 (73%) V-MAGs were not given order-level taxonomy because of the limited genomic data of environmental viruses in the reference database.

### Metaepigenomic analysis

A total of 178 and 42 candidate modified motifs were detected in 108 (46%) P-MAGs and 15 (9%) V-MAGs, respectively (Data S2). Mapped subread coverages of the modified motifs were compatible with P-MAGs and V-MAGs that ranged from 30.6 to 508.9× and 88.3 to 568.8×, respectively. The detected motifs were composed of 59 unique motifs, including 32 motifs with palindromic sequences that allow double-strand modification. Among the unique motifs, 27 and 23 were classified as m6A and m4C methylation types, respectively. Although the current SMRT sequencing technology does not support the detection of the m5C motif, we found four candidate m5C motifs with high subread mean coverages (259× on average). Among the methylated motifs from P-MAGs, 57 (35%) showed <50% modification ratios on the genome, possibly because of the weak detection power of modification from subreads or the existence of strain-level epigenomic heterogeneity in the microbial communities. The modification types of the other 5 motifs were unclassified and possibly represented chemical modifications out of the above three methylations, such as phosphorothioation (*24*). The unclassified motifs showed low modification ratios (ranging from 14–45% with 30% on average), similar to previous observations of phosphorothioated motifs in *Escherichia coli* (12%) and Thaumarchaeota (20%) strains (*22, 23*).

Among the P-MAGs with methylated motifs, G**<u>A</u>**TC was detected most frequently (41 P-MAGs), followed by G**<u>A</u>**NTC (28 P-MAGs), **<u>C</u>**GCG (19 P-MAGs), and BAAA**<u>A</u>** (9 P-MAGs), where B=C/G/T and N=A/C/G/T, and the underlined boldface indicates methylation sites. Among the V-MAGs, RG**<u>C</u>**Y (9 V-MAGs) was the most abundant motif, followed by **<u>C</u>**CNGG (4 V-MAGs), GGW**<u>C</u>**C (3 V-MAGs), and GGH**<u>C</u>**C (3 V-MAGs), where R=A/G, Y=C/T, W=A/T, and H=A/C/T. It is worth noting that, even considering some vague motifs, at least 15 motifs (i.e., BAAA**<u>A</u>**, ACAA**<u>A</u>**, CAA**<u>A</u>**T, CT**<u>A</u>**G, G**<u>A</u>**TGG, G**<u>A</u>**TCC, GTN**<u>A</u>**C, GTW**<u>A</u>**C, S**<u>A</u>**TC, TGNC**<u>A</u>**, TS**<u>A</u>**C, **<u>C</u>**TCC (m4C), GCG**<u>C</u>** (m4C), GGW**<u>C</u>**C (m4C), and TGG**<u>C</u>**CA (m5C), where S=C/G) did not match the known recognition sequences of MTases in REBASE repository. In addition, motifs likely catalyzed by Type I MTases, which are generally characterized as bipartite sequences with a gap of unspecified nucleotides (e.g., **<u>A</u>**TGNNNNNTAC), were undetected in all the P-MAGs and V-MAGs. This result indicates that Type I RM systems were scarce in the epipelagic and mesopelagic prokaryotes and viruses. Regarding vertical distribution of the modified motifs along with the water column, no clear relationship between the frequency of the motif and their habitats was observed; 0.65, 1.2, 0.76, and 0.84 motifs were detected on average in CM1_5m, Ct9H_90m, CM1.5m_200m, and Ct9H_300m P-MAGs, respectively.

### Prediction of MTases and corresponding methylated motifs

To identify MTases that catalyze methylation of the detected motifs, systematic annotation of MTase genes was performed. Sequence similarity searches against known genes stored in REBASE (*43*) identified 171, 43, and 7 of M, R, and RM genes, respectively, from 112 (48%) P-MAGs (sequence identities ranged 20–92%) (Data S3). Among the M and RM genes from P-MAGs, m6A (64%) was the most abundant MTase type, followed by m4C (14%) and m5C (10%), as found in the HiFi read analysis (Fig. 1c). Among the MTase types, Type II MTases were the most abundant (82%), while 9% and 6% genes showed the highest sequence similarity to Type I and III MTases, respectively. This trend is also consistent with the HiFi reads analysis that Type I and III MTases were scarcely detected in the communities (Fig. 1b). Only three genes encoded DNA sequence-recognition proteins, known as the S subunit in the Type I RM system. Most of the MTases were orphan and only four pairs of Type II MTase and REase gene were predicted to possess the same motif sequence specificity and were adjusted on the genome, which may constitute intact Type II RM systems. Other known antiviral defense systems associated with DNA modification—BREX (*45*) and DISARM (*46*) were surveyed; however, no MTase genes likely associated with these systems were found through the P-MAGs. Overall, our analyses highlighted the previously unknown diverse MTases in epipelagic and mesopelagic prokaryotic communities and suggest that the methylation systems play unexplored roles apart from the known defense mechanisms of exogenous DNA.

A total of 58 (20%) MTase genes from P-MAGs showed the best sequence similarity to MTases, whose specificity was exactly matched to the motif identified in our metaepigenomic analysis (Data S2, S3). For example, CM1_200m.P15 contained one MTase that showed the best sequence similarities to those that recognized C**<u>C</u>**SGG, which was perfectly congruent with the motif detected from P-MAG. For CM1_200m.P39, two MTases similar to those that recognize either TTA**<u>A</u>** or **<u>C</u>**GCG were identified, and these motifs were congruently detected in the genome. In Ct9H_300m.P17, five MTases were predicted, two of which were similar to the known MTases that recognize either AG**<u>C</u>**T or G**<u>A</u>**TC, and all of the detected methylated motifs in the genome were completely matched, suggesting that the two MTases were active while the other three were inactive.

In contrast, at least one methylated motif was detected in 40 (17%) P-MAGs, while no MTase gene was found. We assumed that the corresponding MTase genes were missed because of insufficient genome completeness or that these MTase genes diverged considerably from known MTase genes. At least one MTase gene was found in 44 (19 %) P-MAGs, but no methylated motifs were detected. We anticipate that the MTase genes were inactive, or the corresponding methylated motif was undetected due to the low sensitivity of SMRT sequencing, especially in m5C modification (*19, 20*).

Among the viral genomes, 82, 13, and 16 of the M, R, and RM genes were identified from 49 (30%) V-MAGs (sequence identities ranged 23–73%) (Data S3). Similar to the case of P-MAGs, Type II MTases were the most abundant (79%), followed by Type I (7%), and no Type III MTase was detected through the V-MAGs. In contrast to P-MAGs, m4C (62%) was the most abundant modification type in V-MAGs, followed by m6A (30%) and m5C (1%). All the MTases and methylated motifs were unmatched in V-MAGs, except for three pairs (GAT**<u>C</u>** in CM1_5m.V34, G**<u>A</u>**TC, and GTNN**<u>A</u>**C in Ct9H_90mV1), possibly because of the very few viral MTases stored in REBASE Gold Standard database, where currently 16 viral MTase genes were found in a total of 1938 MTase genes.

### Exploration and experimental verification of MTases with new methylated motifs

Among the detected MTase genes, 132 (74%) and 94 (96%) MTases from P-MAGs and V-MAGs, respectively, showed inconsistency between the recognition motifs of their closest relatives and the methylated motifs identified in our metaepigenomic analysis (Data S2, S3). The result suggested that the homology-based estimation of MTase specificity was not sufficient, as in our previous metaepigenomic study of the freshwater microbiome (*29*). To reveal the catalytic specificity of these MTases, we investigated potential pairs of MTase and methylated motif as follows: 1) MTase and methylated motifs were present in the same genome and novel correspondence was estimated; 2) modification types (i.e., m4C, m5C, and m6A) of MTase and methylated motifs were concordant, and 3) the complete sequence of the MTase gene was retrieved. Subsequently, methylation specificities of the selected MTases were experimentally verified with heterologous expression in *E. coli* (Data S4). Briefly, plasmids with one artificially synthesized MTase gene were constructed and transformed into *E. coli* cells, and the methylation status of the isolated plasmid DNA was subsequently observed by REase digestion after heterologous expression. Viral MTases were not selected for the experiment to improve the efficiency of heterologous expression.

In Actinobacteria, Ct9H_300m.P26, one m6A MTase gene, and two m6A and m4C motifs were detected, but none of the MTase and motif matched each other. Thus, we predicted that Ct9H300mP26_1870, whose closest homolog encoded an MTase that exhibits CTCG**<u>A</u>**G methylation activity, would encode an MTase that recognizes BAAA**<u>A</u>**, whereas the motif sequence was not registered in REBASE and no MTase was currently reported to recognize the motif. The REase digestion assay was consistent with the hypothesis that ScaI (AGTACT specificity) did not cleave the BAAA**<u>A</u>**GTACT sequence, which overlapped with BAAAA and AGTACT sequences, on the plasmids only when MTase was expressed in the cells (Fig. 2a). We named this protein M.AspCt9H300mP26I, as a novel MTase that possesses BAAA**<u>A</u>** specificity.

**Fig. 2.**
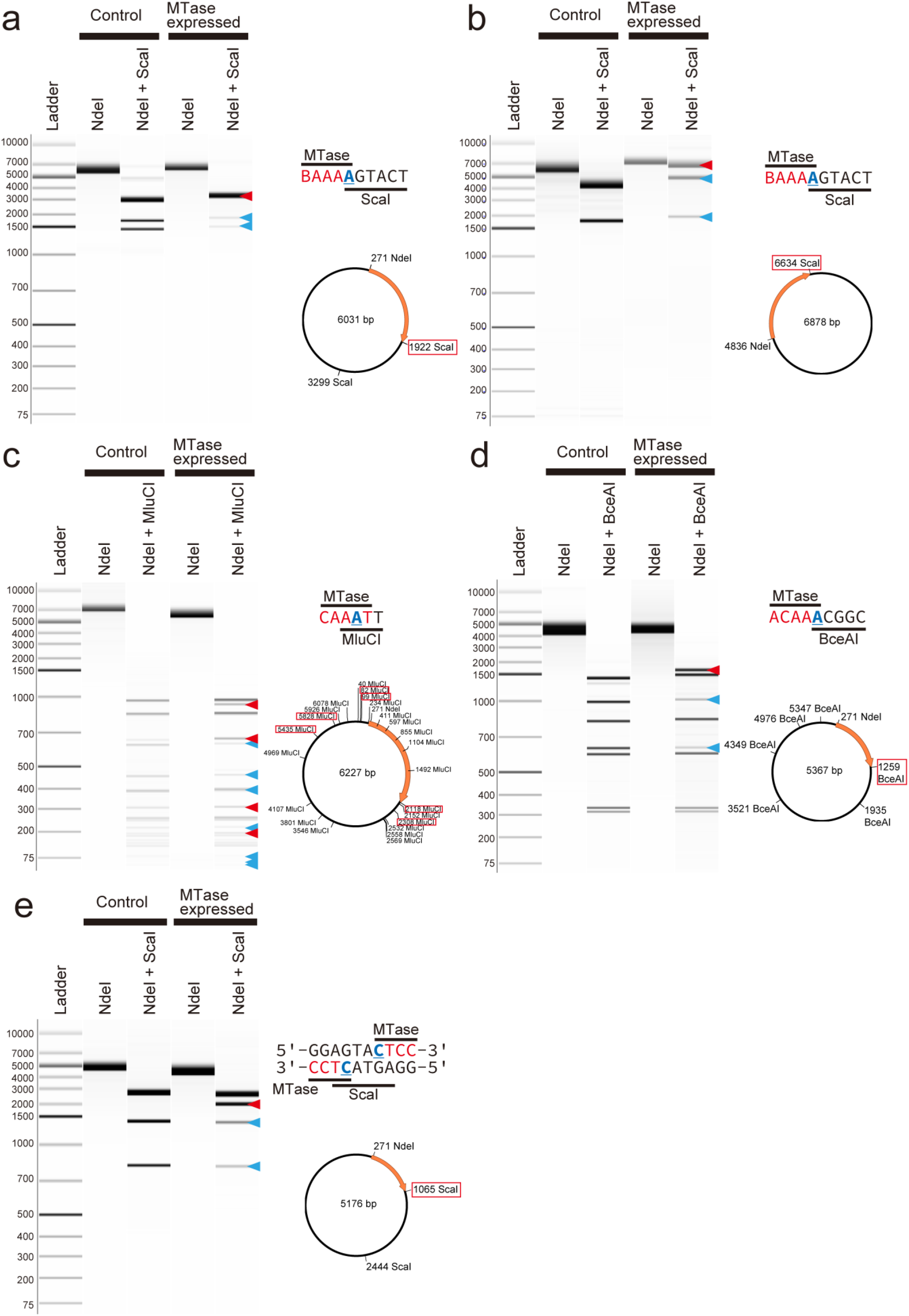
REase digestion assays of MTases with novel specificity. **a** Assay of the Ct9H300mP26_1870 gene. ScaI was used, where the plasmid contained two AGTACT target sites. Within the two sites, one of the target sites was BAAAAGTACT, where overlapped BAAAA and AGTACT were recognized by the MTase and REase. The schematic representation and plasmid map are presented on the right side. The orange arrow represents the transferred gene, and the digestion sites with a red frame represent the location of the overlapped sequence. **b** Assay of the Ct9H90mP5_10800 gene. ScaI was used, where the plasmid contained one AGTACT target site in the BAAAAGTACT site. **c** Assay of the CM1200mP2_32760 gene. MluCI were used, where the plasmid contained 23 AATT target sites. Within them, the six target sites were CAAATT, where overlapped CAAAT and AATT were recognized by the MTase and REase, respectively. **d** Assay of the CM15mP129_7780 gene. BceI were used, where the plasmid contained six ACGGC target sites. Within them, one of the target sites was ACAAACGGC, where overlapped ACAAA and ACGGC were recognized by the MTase and REase, respectively. **e** Assay of the CM1200mP10_13750 gene. ScaI were used, where the plasmid contained two TCATGA target sites. Within them, one of the target sites was GGAGTACTCC, where a pair of CTCC and GGAG (comprehensive sequence of CTCC) and TCATGA were recognized by the MTase and REase, respectively. The pCold III (**a**,**c**-**e**) and pET-47b(+) (**e**) were used as expression vectors. The band sizes were logically expected to emerge (red triangles) and reduce (blue triangles) when the induced MTase causes methylation. All plasmid DNAs were linearized using NdeI.

In Actinobacteria, Ct9H_90m.P5, two MTase genes, and three methylated motifs were detected, and a pair of MTase and motif was concordantly matched, whereas the other MTases did not match any motifs. The latter MTase gene Ct9H90mP5_10800 showed moderate sequence similarity (32%) with a low E-value (1e-70) to M.AspCt9H300mP26I using BLASTP search, and either of the remaining motifs was m6A and m4C. Thus, we predicted that Ct9H90mP5_10800 MTase, whose closest homolog is an m6A MTase that exhibits ATTA**<u>A</u>**T methylation, would have BAAA**<u>A</u>** specificity. As expected, the REase digestion assay showed that ScaI did not cleave the BAAA**<u>A</u>**GTACT sequence on the plasmids only when the protein was expressed (Fig. 2b). Thus, we named this protein M.AspCt9H90mP5I, as a novel MTase that possesses BAAA**<u>A</u>** specificity. We should note that another candidate MTase gene, Ct9H90mP30_5500, detected in Actinobacteria Ct9H_90m.P30 was estimated to possess the same BAAA**<u>A</u>** specificity and showed moderate (33%) and high (87%) sequence similarities to M.AspCt9H300mP26I and M.AspCt9H90mP5I, respectively, although the protein was insolubilized in *E. coli*, resulting in no clear cleavage inhibition in our experiment.

A Planctomycetes CM1_200m.P2 had three MTase genes and two methylated motifs. One of the MTases showed the highest sequence similarity to those recognizing TTA**<u>A</u>** with high similarity (64%). The other CM1200mP2_32760 and CM1200mP2_5150 MTases showed the highest sequence similarity to those catalyzing m6A modification and recognizing GTTA**<u>A</u>**C and ATTA**<u>A</u>**T, respectively, with low similarity (37% and 25%, respectively). The two detected motifs were GCG**<u>C</u>** (m4C) and CAA**<u>A</u>**T (m6A), the latter of which was not found in REBASE. Thus, we expected that either or both MTases would recognize and methylate the novel CAA**<u>A</u>**T motif. The construct CM1200mP2_32760 was not successfully prepared in our experiment, likely because the protein was toxic to *E. coli*. In contrast, CM1200mP2_5150 MTase showed that MluCI (AATT specificity) did not cleave all sequence sites CAA**<u>A</u>**TT on the plasmids only when MTase was expressed, clearly indicating that MTase recognizes CAA**<u>A</u>**T (Fig. 2c). Accordingly, we named the protein M.PspCM1200mP2I as a novel MTase that possesses a previously unknown CAA**<u>A</u>**T specificity.

Chloroflexi CM1_5m.P129 had one MTase gene, which showed the highest sequence similarity to those recognizing TCTAGA (whose modification type and position were not reported). However, the only methylated motif detected in the genome was ACAA**<u>A</u>**, which no MTase was currently reported to recognize. Thus, we hypothesized that CM15mP129_7780 MTase should recognize and modify this novel motif. The REase digestion assay was consistent with the hypothesis that BceAI (ACGGC specificity) did not cleave the sequence site ACAA**<u>A</u>**CGCG only when MTase was expressed (Fig. 2d). Accordingly, we named this protein M.CspCM15mP129I, as a novel MTase that possesses previously unreported ACAA**<u>A</u>** specificity.

In *Ca*. Marinimicrobia CM1_200m.P10, one MTase gene, and one methylated motif were detected. The reported recognition motif of the closest MTase is GAAGA (the modified base is the second position of the complementary sequence T**<u>C</u>**TTC), while the detected motif was **<u>C</u>**TCC. Thus, we hypothesized that the recognition motif of CM1200mP10_13750 MTase would be **<u>C</u>**TCC, a previously unreported methylated motif. The REase digestion assay showed that ScaI was inhibited to cleave the GGAGTACTCC sequence site, where the ScaI targeting site was complementally sandwiched by **<u>C</u>**TCC (Fig. 2e). We accordingly named this protein M.MspCM1200mP10I.

Furthermore, we conducted a re-sequencing analysis to examine the methylation status of the chromosomal DNA of *E. coli* that each novel MTase gene was transformed and expressed. As a result, two BAAA**<u>A</u>**, CAA**<u>A</u>**T, ACAA**<u>A</u>**, and **<u>C</u>**TCC were successfully recalled in the genomes of *E. coli* (Table S4).

### Phylogenetic distribution of modified motifs

To investigate the phylogenetic distribution of the DNA modification system in the MAGs, we used 117 P-MAGs (>20% completeness) and all 163 V-MAGs for robust phylogenetic tree reconstruction, and visualized the modification ratios of the detected motifs in each genome (Fig. 3). Within the P-MAGs, modified motifs were sporadically distributed across the phyla, whereas some showed great concordance with the phylogenetic clades. For example, within the phylum Actinobacteria, **<u>C</u>**GCG and BAAA**<u>A</u>** were spread in all genomes of the class Acidimicrobiia but were not detected in the class Actinobacteria. In contrast, A**<u>A</u>**TT was found in three P-MAGs belonging to a subclade in Acidimicrobiia. TTA**<u>A</u>** was found in four P-MAGs in Chloroflexi. G**<u>A</u>**TC was detected with moderate to high modification ratios (19–99%) through archaeal P-MAGs with two exceptions; no significant G**<u>A</u>**TC signature was detected in Euryarchaeota Ct9H_90m.P24 (7%) and CM1_5m.P82 (0.4%) possibly due to the weak or absent methylation activity in the host organisms. AG**<u>C</u>**T was observed in all two Thaumarchaeota P-MAGs with high modification ratios (82–91%). **<u>C</u>**GCG was found in members from three phyla across the domain: Actinobacteria, Chloroflexi, and Euryarchaeota. We also found that G**<u>A</u>**NTC/G**<u>A</u>**WTC appeared in all 26 Alphaproteobacteria P-MAGs with only one exception, indicating great conservation of the methylation systems in the group as described below. Other than the methylation, AGCT modified motif showed weak modification ratios (2–19%) through the class *Ca*. Poseidoniia P-MAGs, although the motif was detected only in Ct9H_300m.P10 in motif prediction analysis. This result emphasizes that the phylogeny-based modification ratio analysis is efficient for analyzing infrequently modified motifs.

**Fig. 3.**
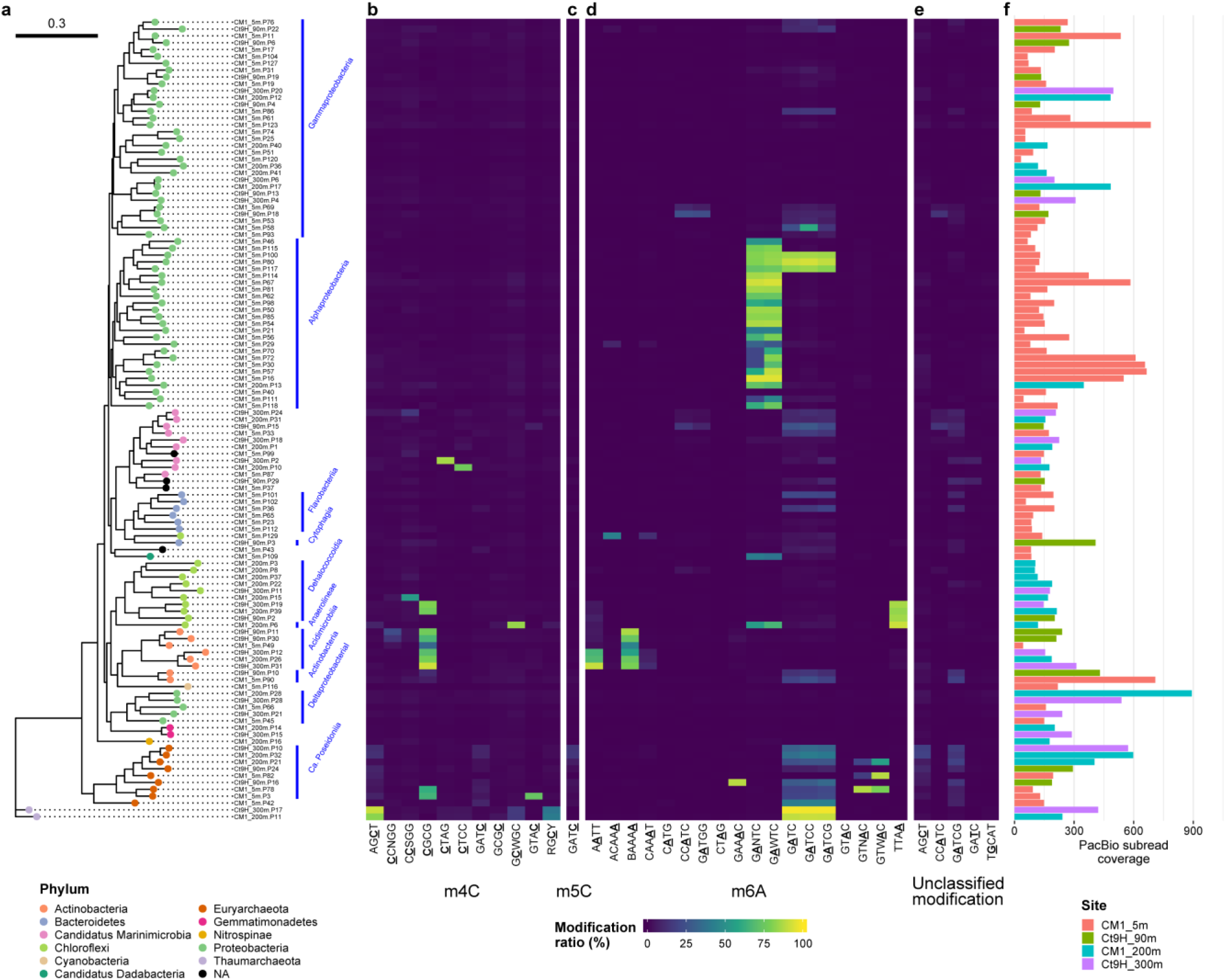
Methylomes of P-MAGs. **a** A phylogenetic tree was constructed using a set of up to 400 conserved bacterial marker genes via the maximum-likelihood method. Node color indicates taxonomy at the phylum level. Nodes were grouped at class to family levels if estimated (blue bars and texts). **b,c,d,e** Modification ratios of detected motifs per genome. **b** m4C, **c** m5C, **d** m6A, and **e** unclassified modifications were individually shown. Motifs detected from P-MAGs without spurious sequence were used. The color range from blue over green to yellow represents modification ratios of motifs on each genome. We should note that modification ratios were affected by overlapped motif sequences; for example, G**<u>A</u>**TCC is completely overlapped by G**<u>A</u>**TC, and both motifs showed similar modification rates in their genomes except in Gammaproteobacteria CM1_5m.P58 where G**<u>A</u>**TCC was detected on the genome from the metaepigenomic analysis and concordantly the modification ratio of G**<u>A</u>**TCC was higher than that of G**<u>A</u>**TC. **f** Coverages of subread on each genome. The bar color represents the source sample of the genome.

In sharp contrast, many motifs showed no clear associations with the phylogenetic topology. For example, G**<u>C</u>**WGC appeared solitary with high modification ratio in Chloroflexi CM1_200m.P6. Similarly, **<u>C</u>**TAG in *Ca*. Marinimicrobia Ct9H_ 300m.P2, **<u>C</u>**TCC in *Ca*. Marinimicrobia CM1_200m.P10, GTA**<u>C</u>** in Euryarcharota CM1_5m.P3, ACAA**<u>A</u>** in Chloroflexi CM1_5m.P129, and GAA**<u>A</u>**C in Euryarcharota Ct9H_90m.P16 were found.

Within all the P-MAGs in this study, no methylated motif was detected in 125 (54%) P-MAGs with high subread coverage (ranging from 31.4–3305.7× and 207.6× on average); thus, this was not addressed by insufficient coverage depth for modification detection. The 125 P-MAGs spread in diverse phyla, such as Proteobacteria, Bacteroidetes, *Ca*. Marinimicrobia, Chroloflexi, Gemmatimonadetes, Cyanobacteria, and Verrucomicrobia. Interestingly, neither methylated motifs nor MTase genes were detected in P-MAGs belonging to several lineages: all two members of Gemmatimonadetes, all two of Nitrospinae, and all five of Verrucomicrobia (Data S1). Methylated motifs were also absent from all three Deltaproteobacteria P-MAGs, although two of them possessed the MTase gene. Within the Gammaproteobacteria P-MAGs, 31 of 52 genomes lacked both methylated motifs and MTase genes. Taken together, these facts suggest the absence of a DNA methylation system in several clades, which is in contrast to a previous study reporting pervasiveness of DNA methylation among bacteria and archaea (*20*). Although further study is required, these observations imply the unexplored benefits of the absence of DNA methylation.

Methylated motifs were occasionally detected with low modification ratios in most V-MAGs, except for Phycodnaviridae and Myoviridae (Fig. 4). Among the Phycodnaviridae V-MAGs, Ct9H_90m.V1 showed G**<u>A</u>**TC and GTNN**<u>A</u>**C with a high modification ratio, whereas Ct9H_90m.V2 harbored TCG**<u>A</u>**. In 14 Myoviridae V-MAGs, 0–5 methylated motifs were detected. However, the proteomic tree showed numbers of V-MAGs that were not assigned but closely related to the Myoviridae family (referred to as ‘Myoviridae-like’). The Myoviridae-like V-MAGs appeared to frequently share several motifs (e.g., RG**<u>C</u>**Y, **<u>C</u>**CWGG, GGW**<u>C</u>**C) with different combinations among them and sometimes harbor additional motifs, while a few numbers of modified motifs were detected in the motif prediction analysis (0.95 motifs per genome on average). This indicates that the taxonomic assignment of the viral genome was frequently missed due to the lack of viral genomes in reference database and the severe underestimation of modified motifs in V-MAGs, likely due to their small genome size (see Materials and Methods). Note that the methylated motifs detected in the V-MAGs were scarcely shared with those in the P-MAGs. No modified motifs other than methylation was found in the V-MAGs. Five Myoviridae-like V-MAGs were predicted to be proviruses, although no clear difference was observed in the modification ratio from the other non-provirus V-MAGs. In 39 Myoviridae-like V-MAGs, several MTases were encoded in their genomes (ranging from 0–8 and 2.4 MTase genes per genome on average), while scarcely detected in the other V-MAGs (ranging from 0–3 and 0.1 on average) (Data S3).

**Fig. 4.**
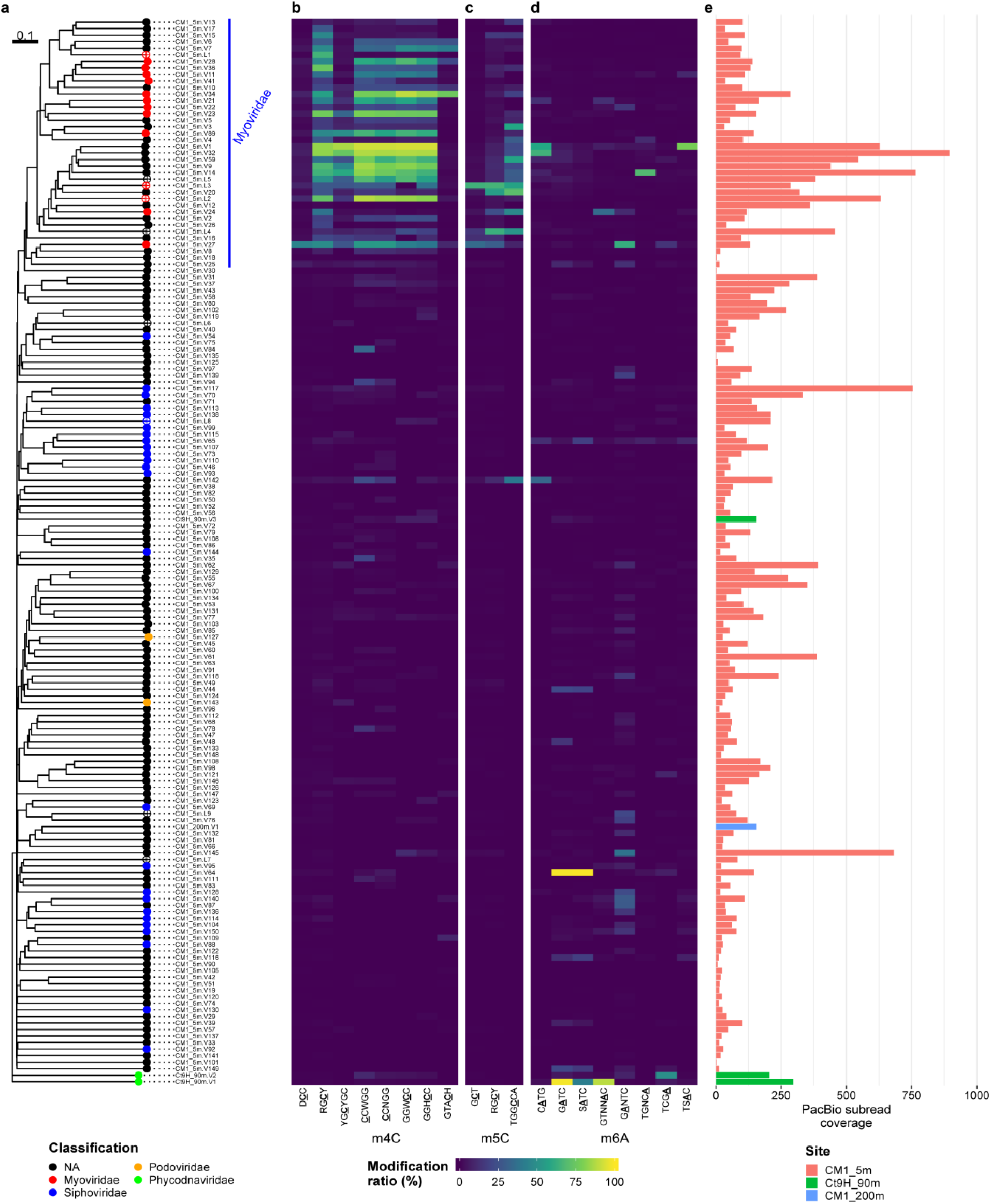
Methylomes of V-MAGs. **a** A proteomic tree was generated based on the global genomic similarities between viral genomes. Proviruses are indicated by circle cross. Node color indicates taxonomy at the family level. **b,c,d** Modification ratios of **b** m4C, **c** m5C, and **d** m6A motifs are presented individually. **e** Coverages of subread on each genome. See Fig. 3.

### MTases that recognize G<u>A</u>DTC/G<u>A</u>WTC motifs in marine Alphaproteobacteria

G**<u>A</u>**NTC methylation is considered to be well conserved in Alphaproteobacteria and is assumed to play an important role in cell cycle regulation via gene regulation (*47*). Indeed, G**<u>A</u>**NTC was previously identified in diverse lineages of Alphaproteobacteria isolates (*20, 48, 49*) and one MAG (*29*) using the modern SMRT sequencing technique, and no alternative motifs have been reported. In our metaepigenomic analysis, G**<u>A</u>**NTC was concordantly detected in 26 of 40 Alphaproteobacteria P-MAGs (Data S3). In addition, we detected similar but different motifs G**<u>A</u>**WTC, G**<u>A</u>**DTC, and G**<u>A</u>**HTC from seven, four, and one Alphaproteobacteria P-MAGs, respectively (where D=A/G/T) (Fig. S6). This result strongly suggests the presence of unknown variations in the methylation system in the lineage. From the Alphaproteobacteria P-MAGs, we predicted 13 complete gene sequences of MTase that were assumed to recognize either of the motifs. However, all of them showed the highest sequence similarity to those known to recognize G**<u>A</u>**NTC with high sequence similarity (47%–80%) (Data S3).

Considering the correspondence of the methylated motifs and MTases, it was expected that four and one MTases would recognize G**<u>A</u>**WTC and G**<u>A</u>**DTC, respectively, rather than G**<u>A</u>**NTC (Data S2, S3). The REase digestion assay of the former four MTases (CM15mP30_3110, CM15mP57_4380, CM15mP70_4410, and CM15mP111_3240) showed that TfiI (GAWTC specificity) cleavage was completely blocked only when MTase was expressed in the cells, whereas HinfI (GANTC specificity) partly cleaved the plasmids (Figs. S7a-d). Despite exhibiting off-target effects under high concentrations of the enzyme, known as ‘star activity’ in REases (*50, 51*), assays of purified CM15mP111_3240 MTase protein suggested that its canonical specificity was G**<u>A</u>**WTC (see Note S1, Figs. S8a-c). The digestion pattern in the assay of CM15mP20_30 was also congruent with the hypothesized G**<u>A</u>**DTC methylation, and re-sequencing analysis successfully recalls the methylated motif (Note S2, Fig. S7e, Table S4). In contrast, as expected, robust G**<u>A</u>**NTC specificity was confirmed in the assay of CM15mP16_9820, which completely inhibited both TfiI and HinfI cleavage (Fig. S7f). Accordingly, we named the four (M.PspCM15mP30I, M.AspCM15mP57I, M.PspCM15mP70I, and M.RspCM15mP111I) and one (M.AspCM15mP20I) proteins as novel MTases that preferentially recognize G**<u>A</u>**WTC and G**<u>A</u>**DTC, respectively, and the last one protein (M.PspCM15mP16I) as G**<u>A</u>**NTC.

M.CcrMI, also known as ‘cell cycle-regulated MTase’ (CcrM) from *Caulobacter crescentus* and with G**<u>A</u>**NTC specificity, is one of the model proteins of prokaryotic MTase (*20, 47, 52*). Based on the sequence alignment of the 13 MTases with M.CcrMI and its homologs, a glycine residue (corresponding to Gly40 in M.CcrMI) was roughly conserved in all MTases with G**<u>A</u>**NTC specificity, while it was replaced with lysine or aspartic acid in all with G**<u>A</u>**WTC (Fig. S9). It has been reported that the M. CcrMI protein contains a substructure that forms a pocket to accommodate the third position of the recognized motif (i.e., nucleotide ‘N’ in G**<u>A</u>**NTC); two hydrophobic residues, Leu38–Leu42 stacks, and flexible Gly39 and Gly40 allow the acceptance of variable nucleotides in the position (*53*). This led to the hypothesis that a replacement of lysine/aspartic acid with glycine at the bottom of the fitting pocket would trigger physical interference in the third position of the motif sequence and change its sequence specificity (i.e., shift from G**<u>A</u>**WTC to G**<u>A</u>**NTC). To test this hypothesis, we constructed a substitution mutant D49G of CM15mP111_3240 (the position corresponding to M.CcrMI Gly40) and conducted the REase digestion assay. However, the mutant showed partial inhibition of HinfI cleavage as compatible with the original MTase, suggesting that another factor rather than merely Gly40 residue defined the third position of the motif as ‘W’ (Fig. S8d).

### Evolutionary history of methylation systems in Alphaproteobacteria

To understand the evolutionary relationships among the MTases recognizing G**<u>A</u>**NTC/G**<u>A</u>**DTC/G**<u>A</u>**WTC motifs in Alphaproteobacteria, we analyzed the phylogenetic diversity of the methylated motifs and the frequencies of the motif sequences on each Alphaproteobacteria P-MAGs (Figs. 5a and c). In Rhodospirillales, SAR116, and Rhodobacteraceae P-MAGs, all four subsets of G**<u>A</u>**NTC (i.e., G**<u>A</u>**ATC, G**<u>A</u>**TTC, G**<u>A</u>**CTC, and G**<u>A</u>**GTC) showed high modification ratios. However, in one Rhizobiales and four SAR11 P-MAGs, G**<u>A</u>**WTC was methylated with higher modification ratios, whereas G**<u>A</u>**STC (i.e., G**<u>A</u>**CTC and G**<u>A</u>**GTC) was almost unmethylated.

**Fig. 5.**
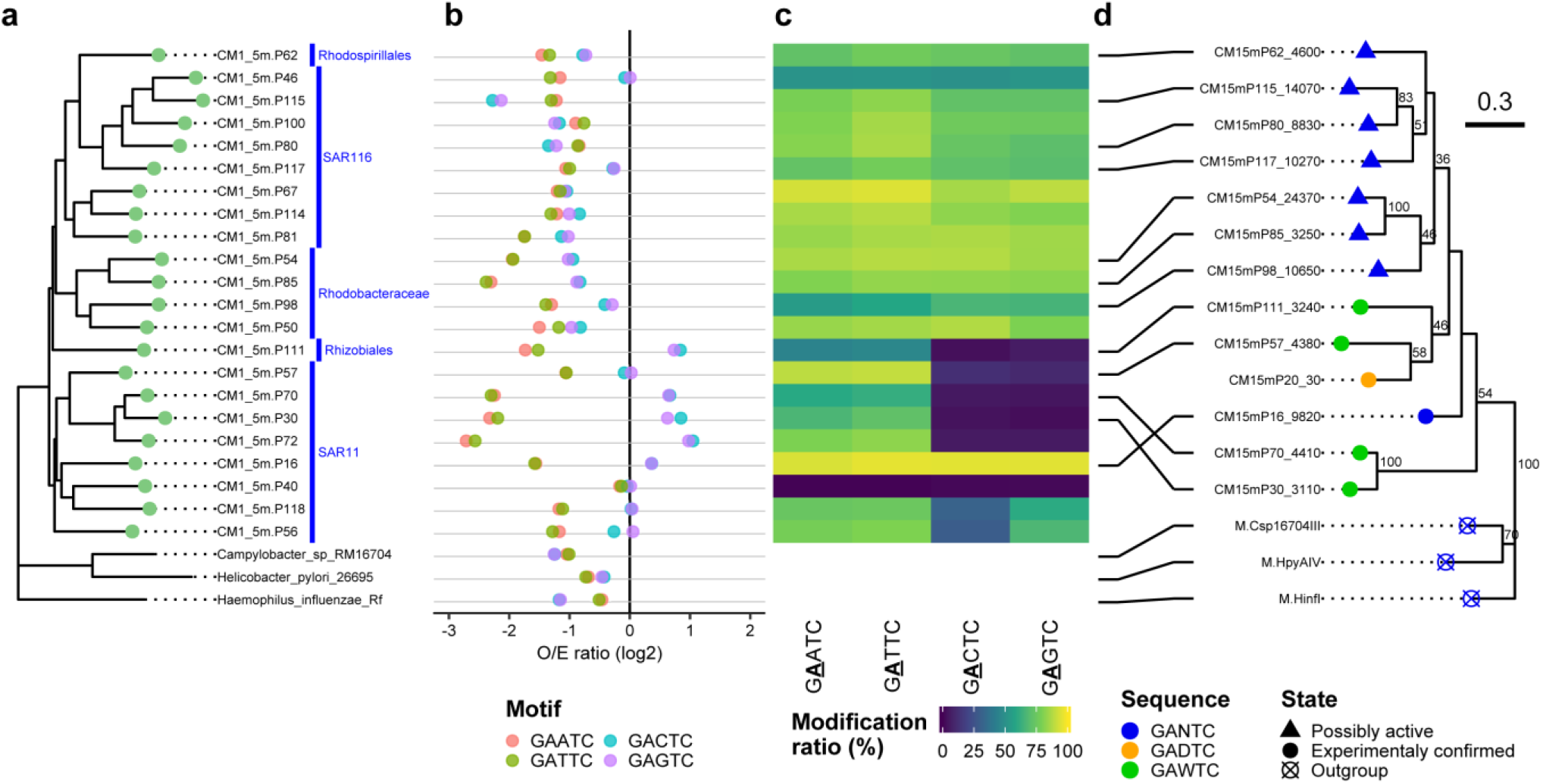
Methylomes and phylogenetic analysis of Alphaproteobacteria P-MAGs. Three homolog MTases, which were found in Proteobacteria isolates and previously confirmed to recognize G**<u>A</u>**NTC were retrieved from REBASE and used as outgroups in this analysis; M.HpyAIV from *Helicobacter pylori* (Epsilonproteobacteria), M.Csp16704III from *Campylobacter* sp. (Epsilonproteobacteria), and M.HinfI from *Haemophilus influenzae* (Gammaproteobacteria). P-MAGs with >25% completeness were used in this analysis for robust phylogenetic tree prediction. **a** A phylogenetic tree of the Alphaproteobacteria P-MAGs. **b** Observed/Expected (O/E) ratio of the GANTC member. A pair of GAATC and GATTC sequences constitutes GAWTC, and all the four sequences (GAATC, GATTC, GACTC, and GAGTC) constituted the GANTC motif, where W=A/T and N=A/C/G/T. **c** Modification ratios of each G**<u>A</u>**NTC component. Blank rows indicate the outgroup whose methylation data was not available. **d** Phylogenetic tree of the MTase genes. Supporting bootstrap values greater than 40% are shown. The node shapes indicate MTases that were estimated (rectangle nodes) or experimentally confirmed (circle) whose specificity and outgroups were indicated by circle cross. The node colors indicate the specificity of each MTase.

The phylogenetic topologies of P-MAGs and MTase were matched in Rhodospirillales, SAR116, Rhodobacteraceae, and Rhizobiales, suggesting good conservation of the MTases in these clades (Figs. 5a and d). In contrast, those in SAR11 showed incongruence with them, possibly because of the weak robustness of the phylogenetic inference of the MTases supported by low bootstrap values. Neither G**<u>A</u>**NTC/G**<u>A</u>**DTC/G**<u>A</u>**WTC methylation nor the corresponding MTase was detected in SAR11 CM1_5m.P40 (Data S2 and S3), indicating that the organism lacked the methylation system. Regardless of the inconsistent topologies between organisms and proteins, MTases likely comprise a monophyletic group. Thus, it is suggested that the methylation systems have been maintained in Alphaproteobacteria and the MTases with G**<u>A</u>**WTC/G**<u>A</u>**DTC specificity branched out from those with G**<u>A</u>**NTC.

Notably, the frequency of motif sequences in the genomes was less than expected when the motif sequences were methylated (Figs. 5b and c). In Rhodospirillales, SAR116, and Rhodobacteraceae P-MAGs, in which G**<u>A</u>**NTC was highly methylated, log2 Observed/Expected ratio (O/E ratio) of all subset of GANTC sequence was −1.14 ± 0.53 (s.d.) on average. This means that GANTC sequences present with >2-fold lower frequency than expected from random distribution on their genomes, suggesting the existence of negative pressure against GANTC sequences. In contrast, in Rhizobiales and SAR11 P-MAGs, except for CM1_5m.P40, the GAWTC O/E ratio was significantly lower than that of GASTC (−1.73 ± 0.59 and 0.43 ± 0.42, respectively) (*p* < 0.05, U-test). The difference suggests a strong negative pressure on the GAWTC sequence, which was attenuated on GASTC. In SAR11 CM1_5m.P40, G**<u>A</u>**NTC was free from methylation and concordantly the O/E ratios were approximately zero (−0.09 ± 0.08), suggesting a weak or no selective pressure under GANTC sequence on the genome.

To gain a more global view of the GANTC sequence representation in the extensive Alphaproteobacteria class, we calculated the O/E ratios using 112 and 195 accessible genomes that covered diverse Alphaproteobacteria (*54*) and all major subclades (I–V) of SAR11 (*44, 55*), respectively (Data S5). All constituent sequences of GANTC generally showed negative O/E ratios in Rhodospirillales, Sphingomonadales, Rhizobiales, Caulobacterales, and Rhodobacterales (−1.78 ± 0.59) (Figs. S10a and b). In contrast, those of Rickettsiales and Holosporales, including numbers of endosymbiotic members, were temperate (−0.39 ± 0.20). Only in SAR11, O/E ratios of GAWTC were significantly lower than those of GASTC (−1.30 ± 0.57 and −0.14 ± 0.37, respectively) (p < 0.05, U-test, Bonferroni correction), and this was concordant with the P-MAG analysis (Fig. 5b). These results indicated that all constituent sequences of GANTC were under negative pressure in Alphaproteobacteria, except Rickettsiales and Holosporales with weak pressure, and SAR11 with selectively attenuated pressure in GASTC constituents. Thus, the O/E ratio profile implied that the G**<u>A</u>**NTC methylation system was not strictly conserved in all Alphaproteobacteria, rather than the G**<u>A</u>**WTC methylation system maintained in the exceptional group.

The estimated phylogenetic tree of SAR11 showed that one and seven P-MAGs belonged to subclades IV and V, respectively (Fig. S10c). The O/E ratios of GAWTC were significantly negative (−1.30 ± 0.45) in contrast to those of GASTC (0.09 ± 0.30) (*p* < 0.05, U-test), but not evenly distributed throughout the SAR11 subclades (Fig. S10d). For example, the GAWTC O/E ratios were higher in subclade Ic (−0.58 ± 0.10), whereas those of GASTC were comparatively lower in subclade Ia.1 (−0.51 ± 0.05). Notably, the GAWTC O/E ratios varied in subclade V (ranging from −2.7 to −0.1). Within subclade V, five minor subclades were identified based on the phylogenetic topology associated with the O/E ratio. One minor subclade, here we referred to Va, showed the lowest GAWTC O/E ratio (−1.83 ±0.59) through the SAR11 subclades. In contrast, subclades Vb, Vc, and Vd showed comparatively higher GAWTC O/E ratios (−0.58 ± 0.28, −0.47 ± 0.06, and - 0.62 ±0.04, respectively). The other subclade, Ve, showed comparatively moderate O/E ratios (−1.00 ±0.42), which is compatible with other major subclades. Despite such variations, overall, the O/E ratio profile suggested that the negative selective pressure in the GAWTC sequence is highly conserved but not or weak in the GASTC sequence through the SAR11 subclades. This may be driven by DNA methylation caused by MTases with G**<u>A</u>**WTC specificity, and the fluctuating pressure among the subclades may be associated with ecological and evolutionary niches, although further investigation is required.

### Metaepigenomics for exploring prokaryotic and viral DNA modification systems in the marine environment

Prokaryotic and viral DNA modification systems should play significant biological functions and have been highlighted; however, little is known about their diversity, ecological role, and evolutionary history, especially in the environmental community. Several studies conducted bisulfite sequencing to investigate prokaryotic m5C modifications using environmental samples (*56, 57*), but other m6A and m4C modifications, which are more popular in prokaryotes, were not investigated. Community-level prokaryotic methylomes have been reported recently (*29, 58, 59*), but the community-level viral methylome has not yet been reported.

The present study has conducted a first metaepigenomic analysis of pelagic microbial communities that dominated members not yet cultured with high complexity and successfully acquired unprecedented DNA modifications. From the HiFi read analysis, which was free from any biases induced in PCR or metagenomic assembly processes, MTase genes were distributed with a similar compositions of modification and MTase types through water depth (5–300 mbsl) (Fig. 1). The reconstructed P-MAGs and V-MAGs possessed a number of modified motifs, including novel ones (Data S2). Subsequent REase assay experiments identified 11 MTases responsible for these reactions, including five with novel specificity (Fig. 2, Fig. S7, and Data S3). The complex evolutionary history of the prokaryotic DNA methylation system is previously reported and considered to have resulted from repeated gene loss, duplication, and horizontal gene transfer, even across phylum or domain levels (*14*) as well as changes in MTases in sequence specificity (*60*). This possibility is supported by the solely presented motifs as well as linage-associated motif conservation from the marine prokaryotic and viral communities in this study (Figs. 3 and 4). Moreover, our analysis provided strong evidence that, in marine Alphaproteobacteria, the methylation systems changed their sequence specificity and affected genomic content through evolutionary history (Fig. 5). Consequently, these results demonstrated that metaepigenomics is effective for comparative methylation analysis within and between microbial populations, as well as modification analysis other than the popular methylations.

Meanwhile, the variance of DNA modifications among lineages has already been applied in bioinformatic applications. For example, an approach of metagenomic binning based on the methylation patterns of assembled contigs has been proposed (*61–63*). However, our results indicate that sets of methylated motifs are frequently shared within phylogenetically close lineages at even higher taxonomic levels such as phylum or order (Fig. 3), and thus could be worthless for distinguishing contigs into individual genome bins.

From the other perspective, careful attention should be paid to the fact that metaepigenomic analysis is based on assembled ‘consensus’ genomes that may overlook epigenomic heterogeneity at lower taxonomic levels such as strain and species. Recent studies have reported possible variations in sets of methylated motifs and MTase genes at the genus to strain levels in wide prokaryotic lineages (*21, 64–66*). Resolving the strain-level diversity of DNA modifications in complex metagenomic samples remains a challenge.

### A possible function of DNA methylation in marine prokaryotic and viral communities

The major detected MTase genes in P-MAGs were orphan and lacked a cognate REase gene, implying that most MTases in pelagic prokaryotes are inactive for protection against extracellular DNA and viral invasion via known physiological systems such as RM (*67*), BREX (*45*), and DISARM (*46*). The similar relative abundance of the MTase genes and their composition in the microbial communities (Fig. 1) suggest that the effects of environmental factors changing with water depth are limited. The possible role of the methylation systems is a factor involved in gene regulation. In addition, because solar UV radiation at the sea surface damages prokaryotic DNA (*68, 69*), some DNA methylations may function for DNA mismatch repair to exhibit photo stress tolerance. In *E. coli*, DNA methylation functions as a marker of the original (parental) DNA strand and facilitates mismatch repair on newly synthesized (daughter) unmethylated strands that frequently occur during cell division under high UV radiation (*4, 5*). It is anticipated that DNA methylation may play a key role in adaptation to the vast marine epipelagic and mesopelagic layers in prokaryotes, although further experimental and proteomic analyses (e.g., transcriptome and metatranscriptome) are required to confirm the epigenetic regulation of the genes involved.

Viruses, the most abundant biological entities in oceanic environments, play diverse roles in marine ecosystems (*70*). Among the V-MAGs, family level variance was found in the methylomes: in Myoviridae V-MAGs, m4C motifs were frequently detected with several m5C and m6A motifs that encode numbers of MTase genes in their genomes; in Phycodnaviridae V-MAGs, several m6A motifs were detected as reported previously (*71*); in the other V-MAGs, including Siphoviridae and Podoviridae, methylation was scarcely detected. These results suggest the existence of strong selective pressure to maintain the methylation system in marine Myoviridae (and possibly Phycodnaviridae), but not in Siphoviridae and Podoviridae. Hence, DNA methylation in Myoviridae (and possibly Phycodnaviridae) may be associated with the genetic roles and ecological strategies of these groups, although the details remain uncertain.

It has been discussed that the main advantageous function of viral MTases is a self-defense weapon against host-encoded defense systems (*25, 72, 73*). However, this hypothesis was not concordant with our results that limited numbers of MTases constituted the known defense systems through the P-MAGs, as discussed above. Thus, the self-defense weapon may play a minor role in DNA methylation in marine viruses. One of the known roles of viral MTase is the initiation of DNA packaging during the late stages of viral infection, found in bacteriophage P1 (*74*). In this system, m6A modification labels ends of the concatemeric viral DNA molecules produced by rolling-circle replication, and the end points, where seven methylated motif sites are clustered in bacteriophage P1, are subsequently cut by an enzyme for DNA packaging into capsids. However, the Myoviridae V-MAGs possessed a variety of m4C motifs with different combinations in their genome (Fig. 4), and the features are likely inefficient to use methylation as a delimiter of concatenated viral DNA replicons. Another possible role of viral methylation is to increase the stability of DNA for dense packing within a viral capsid, as well as alpha-putrescinylthymine modification in bacteriophage φW-14 (*75*). In addition, several viral genes are known to be transcriptionally controlled by a self-encoded MTase originally found in bacteriophage P1 (*76*). The possibility that viral MTase regulates host gene expression to facilitate viral genome replication cannot be ruled out. It would be interesting to explore the role of DNA modification in the viral life cycle.

### Evolutionary history of methylation systems in Alphaproteobacteria

M.CcrMI is one of the most studied MTases to date. M.CcrMI plays significant biological roles such as cell cycle master regulation, and its homologs are highly conserved in Alphaproteobacteria (*47, 48*).All of the known homologs in Alphaproteobacteria are believed to recognize G**<u>A</u>**NTC, and the methylation system was acquired early in the evolution of Alphaproteobacteria (*77*). However, we found unprecedented homologs that possess G**<u>A</u>**WTC and G**<u>A</u>**DTC specificities from members belonging to Rhizobiales and SAR11, and the specificities of these representatives were experimentally demonstrated (Notes S1 and S2, Figs. S6 and S7). Although the phylogenetic topology of the MTases does not match that of the SAR11 subclades in the genomic tree, the MTases with G**<u>A</u>**WTC or G**<u>A</u>**DTC specificity are phylogenetically placed within those of G**<u>A</u>**NTC with high sequence similarity. Thus, all MTases likely comprise a monophyletic group and share a common ancestor rather than acquired from distant lineages by horizontal gene transfer. In addition, it is assumed that the methylation systems have significant importance in the cell cycle process as M.CcrMI, and thus have been under strong selective pressure for maintenance in the orders. The MTase with G**<u>A</u>**WTC and G**<u>A</u>**DTC specificity showed noncanonical G**<u>A</u>**NTC specificity under unoptimized conditions as star activity (Notes S1 and S2, Fig. S7). This enzymatic feature resulted in a scenario in which these protein groups evolved from the ancestral MTase with G**<u>A</u>**NTC specificity by depressing the affinity with GASTC or GACTC sequences. The assay of a mutant protein in which we changed one residue at the bottom of the pocket, which likely accommodated the third position of the motif and distinguished G**<u>A</u>**NTC and G**<u>A</u>**WTC sequences (Fig. S9) showed no obvious specificity shift from G**<u>A</u>**WTC to G**<u>A</u>**NTC (Figs. S8c and d). This result suggests that the GASTC affinity is limited by other or additional residues, or the MTase with G**<u>A</u>**WTC specificity forms a structure distant from M.CcrMI (*53*); thus, further consideration is required.

The O/E ratio analysis showed an oppressive profile of GAWTC sequence present on the genome compared with GASTC in the Rhizobiales and SAR11 P-MAGs. In contrast, GANTC were generally oppressive in the other Alphaproteobacteria P-MAGs, and these were concordant with the detected methylated motifs and specificity of the MTases (Fig. 5). This selection pressure suggests a significant (and may be harmful) effect of methylation on biological processes such as gene expression (i.e., a critical regulatory change driven by methylation at the internal gene coding sequence and/or promoter region). The low frequency of the GANTC sequence in Alphaproteobacteria genomes was previously reported; however, SAR11 (formerly classified in Rickettsiales) was not recognized as an individual group and the frequency of the GAWTC sequence was not evaluated (*48*). Here, our analysis drastically expands the knowledge about methylation in the class that at least a part of SAR11 members possesses the G**<u>A</u>**WTC methylation system. Furthermore, the O/E ratio analysis showed strong and specific negative pressure on the GAWTC sequence in their genomes, in contrast to the other Alphaproteobacteria orders (Fig. S10). Variance in GAWTC O/E ratios among SAR11 subclades showed that different methylation states were associated with the evolution of each subclade. Consequently, our findings provide novel insights into prokaryotic epigenomics that DNA methylation plays a greater role in host evolution than previously recognized. Despite the many challenges of culturing in marine prokaryotes, further investigations are required to evaluate the relationships among molecular function, ecological benefit, and evolution of M.CcrMI homologs in Alphaproteobacteria.

### Effects and challenges of metaepigenomics on environmental microbiology

Our metaepigenomic approach owing to recent improvements in SMRT sequencing illuminates prokaryotic and viral DNA modifications in diverse and complex pelagic microbial ecosystems. It is envisioned that metaepigenomics of prokaryotes and viruses under different ecological niches (e.g., sea area and water depth) and subjects (e.g., soil, gut, and symbionts) will significantly deepen our understanding of the effects of DNA modification and lead to various applications.

Because of the current sequencing read length and throughput of the PacBio platform, it is still challenging to reconstruct genomes of rare lineages, especially in complex microbial communities. Indeed, the HiFi reads covered only half of the microbial communities in this study (Fig. S3), and further sequencing efforts are required for higher-resolution analysis to reveal a whole picture of epigenome in environmental microbial communities. Despite the advantage of the hybrid approach of SMRT and ONT sequencing, each of them requires more than 10 μg of DNA as initial input for library preparation, which allows us to conduct ONT sequencing only for a sample from the surface layer (CM1_5m). In addition, even using current SMRT sequencing technology, only a limited number of DNA modification types can be detected and classified with sufficient reliability (i.e., m4C and m6A), although a number of modifications occur in nature (*2*). Further development of sequencing technology, accurate assembly tools, and reliable modification detection methods will be required for deeper evaluation of prokaryotic and viral DNA methylation in environments.

## Methods

### Seawater sampling

Seawater samples were collected at two close pelagic stations of Japan Agency for Marine-Earth Science and Technology (JAMSTEC) in the northwest Pacific Ocean during JAMSTEC KM19-07 cruises of the *Research Vessel (R/V) Kaimei* in September 2019 (Fig. S1, Table S1). The sampling stations were approximately 180 and 140 km offshore from the main island of Japan, and 60 km from each other. Each 90–300 L of seawater was collected from 5 and 200 mbsl at station CM1 (34.2607 N 142.0203 E) and 90 and 300 mbsl at station Ct9H (34.3317 N 141.4143 E) (referred to as CM1_5m, CM1_200m, Ct9H_90m, and Ct9H_300m, respectively). Sampling permit for expeditions in Japan’s exclusive economic zone was not required as in domestic areas and did not involve endangered or protected species. Seawater from 5 mbsl was directly sampled using a built-in pumping system from the bottom of the ship via approximately 5 m of intake pipe that was designed for continuous monitoring of sea surface hydrography. The valve of the pumping system was opened at least 30 min before starting the sampling to entirely flush the internal water and rinse the pipe. Seawaters from 90, 200, and 300 mbsl were sampled using 12-L Niskin-X bottles (General Oceanic, Miami, Florida, USA) in a CTD rosette system. The vertical profiles of temperature, salinity, and pressure data were obtained using the SBE9plus CTD system (Sea-Bird Scientific, Bellevue, Washington, USA). The vertical profiles of dissolved oxygen (DO) concentrations were obtained using an *in situ* DO sensor RINKO-III (JFE Advantech, Hyogo, Japan) connected to the CTD. The vertical profiles of chlorophyll *a* concentrations were obtained using an *in situ* Fluorometer RINKO profiler (JFE Advantech). The seawater samples in the Niskin-X bottles were transferred to sterilized 20 L plastic bags and immediately stored at 4 °C until further filtration. Filtration was performed with 0.22-μm Durapore membrane filters (Merck KGaA, Darmstadt, Germany) after prefiltration with 5 μm Durapore membrane filters (Merck KGaA) onboard. The filters were then immediately stored at temperatures lower than −30 °C.

### Flow cytometric assessments of prokaryotic cell and viral-like particle abundances

Seawater samples for flow cytometric assessments of prokaryotic cell and viral-like particle (VLP) abundances were obtained every 10–50 m at station CM1 and 10–100 m at station Ct9H, fixed with 0.5% (w/v) glutaraldehyde (final concentration) in 2 mL cryo-vials on board and stored at −80 °C until further analysis. For assessment of prokaryotic cell abundance, 200 μL of each sample was stained with SYBR Green I Nucleic Acid Gel Stain (Thermo Fisher Scientific, Waltham, Massachusetts, USA) (×5 of manufacturer’s stock, final concentration) at room temperature for >10 min. For assessment of VLP abundance, 20 μL of each fixed sample was diluted 10 times with TE buffer and stained with SYBR Green I (×0.5 of manufacturer’s stock, final concentration) for 10 min at 80 °C. Total prokaryotic cells and VLP abundance in 100 μL samples were determined using an Attune NxT Acoustic Focusing Flow Cytometer (ThermoFisher Scientific) by their signature in a plot of green fluorescence versus side scatter (*78, 79*).

### DNA extraction and shotgun sequencing

Microbial DNA was retrieved using a DNeasy PowerSoil Pro Kit (QIAGEN, Hilden, Germany) according to the supplier’s protocol. The filters were cut into 3-mm fragments and directly suspended in the extraction solution from the kit for cell lysis. SMRT sequencing was conducted using a PacBio Sequel system (Pacific Biosciences of California, Menlo Park, California, USA) at the National Institute of Genetics (NIG), Japan. SMRT libraries for HiFi read via CCS mode were prepared with a 5-kb insertion length. Briefly, 4–6 kb DNA fragments from each genomic DNA sample were extracted using the BluePippin DNA size selection system (Sage Science, Beverly, Massachusetts, USA). The SMRT sequencing library of CM1_5m and the other three samples were prepared using the SMRTbell Template Prep Kit 1.0-SPv3 and SMRTbell Express Template Prep Kit 2.0, respectively, according to the manufacturer’s protocol (Pacific Biosciences of California). The final SMRT libraries were sequenced using four, three, three, and three Sequel SMRT Cell 1M v3 for CM1_5m, CM1_200m, Ct9H_90m, and Ct9H_300m, respectively. Nanopore sequencing of CM1_5m was conducted using a GridION Mk1 platform with five flow cells according to the manufacturer’s standard protocols at NIG. ONT libraries were prepared and purified simultaneously by filtering out a small number of fragments using AMPure XP beads (Agencourt BioSciences, Beverly, Massachusetts, USA). Illumina sequencing (2× 300 bp paired-end reads) was conducted using an Illumina MiSeq platform (Illumina, San Diego, California, USA) at JAMSTEC. Illumina libraries were prepared using the KAPA Hyper Prep Kit (Roche, Basel, Switzerland) and mixed with Illumina PhiX control libraries, as described previously (*80*).

### Bioinformatic analysis of sequencing reads and assembled genomes

CCS reads that contained at least five full-pass subreads on each polymerase read and with >99% average base-call accuracy were retained as HiFi reads using the standard PacBio SMRT software package with the default settings. Metagenomic coverage of HiFi reads was estimated using Nonpareil3 with default settings (*81*). For taxonomic assignment of HiFi reads, Kaiju (*32*) in Greedy-5 mode (‘-a greedy -e 5’ setting) with NCBI nr (*33*) and GORG-Tropics databases (*34*) were used. HiFi reads that potentially encoded 16S ribosomal RNA (rRNA) genes were extracted using SortMeRNA (*82*) with default settings, and full-length 16S rRNA gene sequences were predicted using RNAmmer (*83*) with default settings. The 16S rRNA gene sequences were taxonomically assigned using BLASTN (*35*) against the SILVA database release 128 (*84*), where the top-hit sequences with e-values ≤1E-15 were retrieved. Coding sequences (CDSs) with >33 aa length in HiFi reads were predicted using Prodigal (*85*) in anonymous mode (‘-p meta’ setting). For Illumina read data, both ends of reads that contained low-quality bases (Phred quality score < 20) and adapter sequences were trimmed using TrimGalore (https://github.com/FelixKrueger/TrimGalore) with default settings. The remaining paired-end reads were merged with at least 10 bp overlap using FLASH (*86*) with default settings.

HiFi and ONT reads were *de novo* assembled using wtdbg2 (Redbean) with the settings for PacBio CCS and ONT reads, respectively, according to the provided instructions (*87*). Assembled contigs from ONT reads were polished using both HiFi and Illumina short reads and HyPo (*88*). For the polishing, HiFi and Illumina reads were mapped on the pre-polished contigs using pbmm2, an official wrapper software for minimap2 (*89*) with CCS reads settings, and Bowtie2 (*90*) with ‘-N 1’setting, respectively.

The assembled contigs were binned using MetaBAT (*91*) based on genome coverage and tetra-nucleotide frequencies as genomic signatures, where the genome coverage was calculated with Illumina reads using Bowtie2 with ‘-N 1’ setting. The quality of bins was assessed using CheckM (*92*), which estimates completeness and contamination based on taxonomic collocation of prokaryotic marker genes with default settings. Bins with <10% contamination were retrieved according to the metagenome-assembled genome (MIMAG) standards (*93*) and defined as prokaryotic metagenome assembled genomes (P-MAGs). We note that the partial genome would be sufficient for detecting DNA modifications and modified motifs; completeness was not considered for P-MAG definition. Sequences of 16S rRNA genes in each P-MAG were retrieved using RNAmmer (*83*) with default settings. The taxonomy of the P-MAGs was estimated based on 16S rRNA gene sequences, CAT (*94*), and Kaiju (*32*). P-MAGs that were not assigned to prokaryotes or assigned but with low reliance (<0.6 supported score) using CAT were excluded from further analysis. CDSs with >33 aa length in each P-MAG were predicted using Prodigal (*85*) with default settings. Functional annotations were achieved through HMMER (*95*) searches against the Pfam database (*96*), with a cutoff e-value of ≤ 1E-5.

For viral sequence collection, the assembled contigs were screened using VirSorter2 (*97*) with default settings. Quality assessment of the retrieved contigs and removal of flanking host regions from integrated proviruses was performed using CheckV (*98*). Contigs assigned to either ‘Complete’ or ‘High-quality’ or ‘Medium-quality’ were defined as viral metagenome assembled genomes (V-MAGs) and used for further analysis. Taxonomy levels lower than kingdom were estimated using CAT (*94*). CDSs were predicted using Prodigal (*85*) in an anonymous mode (‘-p meta’ setting). Functional annotations were achieved in the same way as for P-MAGs.

### Bioinformatic analysis of modification systems

DNA modification detection and motif analysis were performed in each MAG independently according to the officially provided tool SMRT Link v8.0. Briefly, subreads were mapped to the assembled contigs using pbmm2, and the interpulse duration ratios were calculated. Candidate motifs with scores higher than the default threshold values were retrieved as modified motifs. Those with infrequent occurrences (<50 and <10 in P-MAGs and V-MAGs, respectively) or very low methylation fractions (<10%) in each MAG were excluded from further analysis. Motifs with several ambiguous sequences that were considered to have occurred by misdetection were manually curated. For example, HBNNNNNNVGGW**<u>C</u>**CNH was detected in CM1_5m.V59, where H=A/C/T, B=A/G/T, V=A/C/G, and W=A/W, but this motif represents palindromic GGW**<u>C</u>**C and the spurious partial sequences of former HBNNNNNNV and latter NH were likely due to incomplete detection of the motif. Notably, we frequently found candidate motifs that showed such ambiguity in V-MAGs. This is likely a result of the weak motif estimation power from small genomes; the low presence of motifs in the genome negatively affected the motif-finding algorithm implemented in the MotifMaker tool, which is based on progressive testing for seeking longer motif sequences using a branch-and-bound search.

Genes encoding DNA methyltransferases (MTases), restriction endonucleases (REases), and DNA sequence-recognition proteins (S subunits) were searched using BLASTP (*35*) against an experimentally confirmed gold-standard dataset from REBASE (*43*) (downloaded on February 9, 2021), with a cutoff e-value of ≤ 1E-5. Sequence specificity information for each hit MTase gene was retrieved from REBASE. The flanking regions of the MTase genes were investigated to search for REase genes and to examine whether they constitute RM systems. The BREX (*45*) and DISARM (*46*) systems were sought based on Pfam domains.

For accurate analysis of methylome diversity, P-MAGs with >20% completeness were used for the phylogenetic analysis. A maximum-likelihood (ML) tree of the MAGs was constructed using PhyloPhlan3 (*99*) on the basis of a set of 400 conserved prokaryotic marker genes (*100*) with ‘--force_nucleotides --diversity high --accurate’ settings. The proteomic tree of V-MAGs was estimated using ViPTreeGen (*101*) with default settings.

For estimation of a robust phylogenetic tree of Alphaproteobacteria P-MAGs, those with higher quality (>25% completeness) were retrieved and used for ML tree reconstruction using PhyloPhlan3 with ‘--force_nucleotides --diversity low --accurate’ settings. To calculate the expected/observed (E/O) ratio of each motif sequence, the expected and observed counts of its presentation on the genome were computed using R’MES (*102*) and SeqKit (*103*), respectively. An ML tree of MTases was constructed using MEGA X (*104*)with LG substitution model with a gamma distribution (LG+G), which was selected based on the Bayesian information criterion (BIC), and 100 bootstrap replicates. Three pairs of the Proteobacteria genome and carried MTase homolog gene were retrieved from the NCBI database and REBASE, respectively, and used for outgroups; pairs of *Campylobacter* sp. RM16704 and M.Csp16704III, *Haemophilus influenzae* Rd KW20 and M.HinfI, and *Helicobacter pylori* 26695 and M.HpyAIV. Multiple sequence alignment was calculated using the MTase sequences in addition to M.CcrMI from *Caulobacter crescentus* CB15 using Clustal Omega (*105*).

For phylogenetic tree analysis of Alphaproteobacteria and SAR11 genomes, a total of 112 and 195 deposited genomes were referred to by Muñoz-Gómez et al. (*54*) and Haro-Moreno et al. (*55*) were retrieved from the NCBI database, respectively (Data S5). For the analysis of Alphaproteobacteria, four Betaproteobacteria and four Gammaproteobacteria genomes were retrieved from the NCBI database and used as outgroups. For the analysis of SAR11, genomes of *Rickettsia felis* URRWXCal2, *Rhodospirillum rubrum* ATCC11170, *Rickettsia bellii* RML369-C, and *Acidiphilium cryptum* JF-5 were retrieved from the NCBI database and used as outgroups. The phylogenetic trees were estimated using PhyloPhlan3 with ‘--force_nucleotides --diversity low --accurate’ settings. Subclades of the SAR11 P-MAGs were inferred based on the topology of the phylogenetic tree according to the previous definition (*55, 106, 107*).

### Experimental verification of MTase activities

To verify MTase specificity, selected MTase genes were artificially synthesized with codon optimization by Thermo Fisher Scientific (Data S4). The genes were cloned into the pCold III expression vector (Takara Bio, Shiga, Japan) using the In-Fusion HD Cloning Kit (Takara Bio). Additional specific sequences were inserted downstream of the termination codon for the methylation assay if an appropriate sequence was absent from the plasmid vector. The constructs were transformed into *E. coli* HST04 *dam^-^/dem^-^* (Takara Bio), which lacks the *dam* and *dcm* MTase genes. In addition, constructs of Ct9H90mP5_10800 and Ct9H90mP30_5500 were alternatively induced into the pET-47b(+) expression vector (Merck KGaA) using the In-Fusion HD Cloning Kit and transformed into *E. coli* BL21 Star (DE3) (Thermo Fisher Scientific) due to severe insolubilization of the expressed protein in the former manner. Soluble protein levels were measured using SDS-PAGE analysis as needed. The *E. coli* strains were cultured in LB broth supplemented with the appropriate antibiotics, ampicillin or kanamycin. MTase expression was induced according to the supplier’s protocol for the expression vector. Plasmid DNA was isolated using the FastGene Plasmid Mini Kit (Nippon Genetics, Tokyo, Japan) or NucleoSpin Plasmid EasyPure Kit (Takara Bio). The REase NdeI was employed for the linearization of plasmid DNAs. Methylation status was assayed simultaneously with linearizing digestion using the appropriate REases. All REases were purchased from New England BioLabs (NEB) (Ipswich, Massachusetts, USA). All digestion reactions were performed at 37 °C for 1 h, except for the simultaneous digestion of HinfI and TfiI at 37 °C for 30 min, followed by 65 °C for 30 min.

We further verified MTases with novel motif specificities (i.e., Ct9H300mP26_1870, Ct9H90mP5_10800, CM1200mP2_32760, CM15mP129_7780, CM1200mP10_13750, and CM15mP20_30) by SMRT sequencing. Chromosomal DNA of *E. coli* HST04 *dam^-^/dcm^-^* strains in which target MTases were transformed were extracted using the DNeasy UltraClean Microbial Kit (QIAGEN) according to the supplier’s protocol after induction of gene expression. Multiplex SMRT sequencing was conducted using PacBio Sequel II (Pacific Biosciences of California) according to the manufacturer’s standard protocols. Briefly, 12–50 kb DNA fragments from each genomic DNA sample were extracted using the BluePippin size selection system (Sage Science) for continuous long read (CLR) sequencing. SMRT sequencing libraries were prepared using the SMRTbell Express Template Prep Kit 2.0 and Barcoded Overhang Adapter Kit 8A, according to the manufacturer’s protocol (Pacific Biosciences of California). All final SMRT libraries were sequenced using a Sequel II SMRT Cell 8M. Methylated motifs were detected using SMRT Link v9.0 against the *E. coli* K-12 MG1655 reference genome (RefSeq NC_000913.2).

For the *in vitro* assay of CM15mP111_3240 MTase and its point mutant, recombinant proteins were purified. N-terminal 6×His-tag fusion MTase and D49G mutant were constructed using PCR and cloned into the pCold III expression vector. *E. coli* cells (HST04 *dam^-^/dcm^-^*) transformed with the constructs were grown at 37 °C for 16 h in 20 mL of medium A (LB medium containing 50 μg/mL of ampicillin) with shaking. The culture was then inoculated into 2 L of medium A in a 5-L flask, incubated at 37 °C for 2–3 h with shaking, and grown until the optical density at A_600_ nm reached 1.0. Then, MTase expression was induced at 15 °C with the addition of 0.1 mM isopropyl β-d-1-thiogalactopyranoside (IPTG), and the cultures were subsequently incubated for 16 h according to the manufacturer’s standard protocol of pCold. *E. coli* cells were lysed by sonication in Buffer A [20 mM HEPES-Na (pH 7.5), 150 mM NaCl, 5% glycerol, 1 mM DTT, and 50 mM imidazole]. The cell lysate was centrifuged at 12,000 rpm for 30 min at 4 °C and passed through a GD/X syringe filter with a 0.45-μm pore size (Cytiva, Marlborough, Massachusetts, USA). The supernatant was subjected to a two-column chromatography using ÄKTA prime chromatography system (Cytiva). The presence of the desired protein was confirmed using SDS-PAGE. The sample was loaded onto a 5-mL HisTrap HP column (Cytiva) at a flow rate of 2 mL/min. The column was then washed with Buffer A. The His-tagged protein was eluted with Buffer B [20 mM HEPES-Na (pH 7.5), 150 mM NaCl, 5% glycerol, 1 mM DTT, and 300 mM imidazole]. The eluted fractions were pooled and diluted 5-fold with Buffer C [20 mM HEPES-Na (pH 7.5), 150 mM NaCl, and 1 mM DTT]. The diluted solution was concentrated to approximately 5 mL using 30 kDa molecular weight cutoff Amicon Ultra centrifugal filters (Merck KGaA), passed through a Millex-GP syringe filter with 0.22 μm pore size (Merck KGaA), and loaded onto a HiLoad 16/600 Superdex 200 pg column (Cytiva) pre-equilibrated with Buffer C. The protein was collected as a single peak and concentrated to 2.5 mg/mL (~50 μM in monomer concentration). The protein was aliquoted, flash-frozen in liquid nitrogen, and stored at −80 °C, or preserved with 50% glycerol at −30 °C until used for further assays.

Purified MTases were used for enzymatic methylation. The substrate unmethylated DNAs were produced using PCR with the pCold III vector transferred CM15mP111_3240 gene as a template to match with the *in vivo* assay of MTase. Methylation reactions were carried out in a reaction buffer [20 mM HEPES-Na pH 7.5, 100 mM NaCl, and 100 μg/mL BSA] with 5 nM substrate DNA and 1 μM purified MTase in solution at 20 °C for 1 h unless specified otherwise. To investigate salt sensitivity, the NaCl concentration was varied from 0–400 mM. To investigate the thermal sensitivity, the reaction temperature was varied from 5–40 °C. For investigations of star activity, MTase and glycerol concentrations were varied between 1 and 15 μM and 0–10% v/v, respectively, and the reaction time was extended to 3 h. The reactions were started with 160 μM SAM (NEB) in solution and stopped by adding guanidinium thiocyanate solution buffer NTI (Takara Bio). After the methylation reaction following DNA purification, the methylation status was assayed using HinfI digestion at 37 °C for 30 min.

## Supporting information

Supplementary Informatin

## Data availability

The raw sequencing data and assembled genomes were deposited in the DDBJ Sequence Read Archive and DDBJ/ENA/GenBank, respectively (Data S6). All data are registered under BioProject ID PRJDB11069 [http://trace.ddbj.nig.ac.jp/BPSearch/bioproject?acc=PRJDB11069].

## Acknowledgments

We would like to thank the captain, crew, and onboard scientists and technicians of the R/V *Kaimei* (JAMSTEC) during KM19-07 cruise. The SMRT and Nanopore sequencing were supported by NIG. We thank Keiko Tanaka, Eiji Tasumi, Akiko Makabe, Minoru Hamana, Masahito Shigemitsu, Hiroshi Uchida, Yusuke Tsukatani, Hidetaka Nomaki, Takeuchi Akinori, Shuhei Ota, Yuya Tada, Mancha Mabaso, Jarishma Gokul, and Thulani Makhalanyane for seawater sampling. We are grateful to Masami Koizumi for technical assistance with cell and viral-like particle counting and flow cytometry experiments, and Fumie Kondo and Miwako Tsuda for their helpful suggestions and support in the molecular experiments. This work was financially supported by the Japan Society for the Promotion of Science (grant numbers JP18K11636, JP19H04246, JP19H05667, JP19H05684, JP19K21203, JP20H02020, and JP20K15444), and the Institute for Fermentation, Osaka (IFO).

## Author contributions

SH conceived and designed the study, performed the sampling, molecular experiments, bioinformatics analyses, and wrote the manuscript. TS performed the sampling, designed and performed the molecular experiments and protein purification, and wrote the manuscript. MH performed the sampling and DNA sequencing using Illumina. AT performed DNA sequencing using PacBio and Nanopore. SK designed the cruise. TY designed and performed sampling and wrote the manuscript. TN wrote the manuscript and supervised the project. All authors read and approved the final manuscript.

