## Supplementary Informatin for "Diverse DNA modification in marine prokaryotic and viral communities"

Supplementary notes S1 to S2

Supplementary references

Supplementary figures S1 to S10.

Supplementary tables S1 to S4.

### Supplementary notes

#### Note S1. *In vitro* assay of MTases that possess G**A**WTC specificity.

The REase digestion assays of MTases detected from Alphaproteobacteria P-MAGs and predicted to recognize the G**A**WTC motif showed notable plasmid cleavage patterns associated with incubation temperature after MTase expression (Figures S7a-d). To better understand the MTases, CM15mP111_3240 was chosen as a representative for further *in vitro* assays as an alternative to *in vivo* methylation. The CM15mP111_3240 gene is 367 aa long with a molecular weight of 42.6 kDa, similar to M.CcrMI (359 aa with 39.7 kDa). Initial purification efforts of the protein with N-terminal His-tag constructed using PCR and protein elution according to the previous studies of M.CcrMI (*1*–*4*); however, it resulted in mild insolubilization, possibly due to the high concentration of imidazole (300 mM). Alternative approaches including on-column tag digestion, which involves cleavage of either N- or C-terminal His-tags with human rhinovirus 3C (HRV 3C) protease cleavage site (Leu-Glu-Val-Leu-Phe-Gln-Gly-Pro) were then conducted, but SDS-PAGE analysis also showed significant insolubilization. Therefore, we used the initial approach accompanied with a five-fold dilution of aliquots with storage buffer immediately after protein elution (50 mM imidazole, final concentration) to avoid insolubilization. This strategy resulted in a significantly higher amount of purified protein with high purity, as confirmed using SDS-PAGE.

CM15mP111_3240 MTase appeared to partially block HinfI cleavage under low salt concentrations (0–150 mM NaCl), as expected when G**A**WTC sites were specifically methylated (Figure S8a). In contrast, substantially no additional inhibition was observed when more than 250 mM NaCl was supplied, indicating that high salt concentrations affect enzymatic activity. The assays of thermal sensitivity showed positive enzymatic activity at 5–35 °C (especially fairly active at 15–25 °C), but not at >35 °C (Figure S8b). These profiles strongly support our hypothesis that MTase recognizes G**A**WTC and causes methylation. The temperature-activity profile was concordant with the marine water temperature at the sampling sites (Figure S1b), suggesting that MTase was thermally optimized in an epipelagic environment. Out of the optimal conditions, we found that HinfI cleavage was further inhibited beyond merely G**A**WTC methylation after MTase reaction with higher enzymatic and glycerol concentrations (Figure S8c). This indicates that abnormal conditions lead to relaxing the sequence recognition of MTase, resulting in the promiscuous extension of its specificity from G**A**WTC to G**A**NTC. This phenomenon was similarly observed in the number of REases known as 'star activity', and has also been reported for various MTases (*5*–*11*). Hence, the remarkable inhibition of HinfI cleavage observed in the *in vitro* assay (Figures S7a-d) could be explained by the artificially induced overexpression of the protein lead star activity, which caused the peculiar methylation block against REase digestion. Taken together, CM15mP111_3240 MTase, as well as the other three homologs (i.e., CM15mP30_3110, CM15mP57_4380, and CM15mP70_4410), likely possess G**A**WTC specificity as a canonical motif and exhibit off-target G**A**NTC methylation under non-optimal conditions such as high enzymatic and glycerol concentrations.

#### Note S2. *In vivo* assay of MTases that possess G**A**DTC specificity

Based on the metaepigenomic analysis, a methylated motif G**A**DTC was detected with a high modification ratio (75.4–81.6%) from four Alphaproteobacteria P-MAGs (i.e., CM1_5m.P124, CM1_5m.P118, CM1_5m.P20, and CM1_5m.P56) (Data S3). However, modification ratio analysis showed not absent but weak G**A**CTC methylation (29.6–35.4%) on the P-MAGs (Figure S6), suggesting that the responsible MTase did not strictly recognize G**A**DTC modification rather than still possess a weak affinity for G**A**CTC. Among the P-MAGs, one MTase gene that showed the highest sequence similarity to those with G**A**NTC specificity was found in CM1_5m.P20. Thus, as described above, we hypothesized that CM15mP20_30 MTase possesses novel G**A**DTC specificity with weak G**A**CTC affinity.

The REase digestion assay of MTase showed that TfiI cleavage was almost completely inhibited at all temperatures tested, indicating that MTase possesses G**A**WTC specificity (Figure S7e). In contrast, the HinfI cleavage showed extra upper bands at 5 °C, which were not observed in other MTases with G**A**WTC specificity under the same conditions (Figures S7a-d), and complete inhibition at >10 °C, similar to the MTase with G**A**NTC specificity (Figure S7f). However, this profile was consistent with the hypothesized G**A**DTC specificity: G**A**GTC hemimethylation (where G**A**GTC is methylated while complementary GACTC is not methylated) is sufficient for blocking HinfI cleavage (*12*). Indeed, re-sequencing analysis of *E. coli* in which the MTase gene was transferred successfully recalled G**A**DTC methylated motifs from the cells incubated at 5 °C, whereas G**A**NTC was detected in the cells incubated at 15 °C (Table S4). Taken together, although the molecular mechanism is still unclear, these results suggest that CM15mP20_30 MTase recognizes G**A**DTC as canonical and possesses weak G**A**CTC specificity, which may be affected by external conditions such as temperature.

#### Supplementary figures

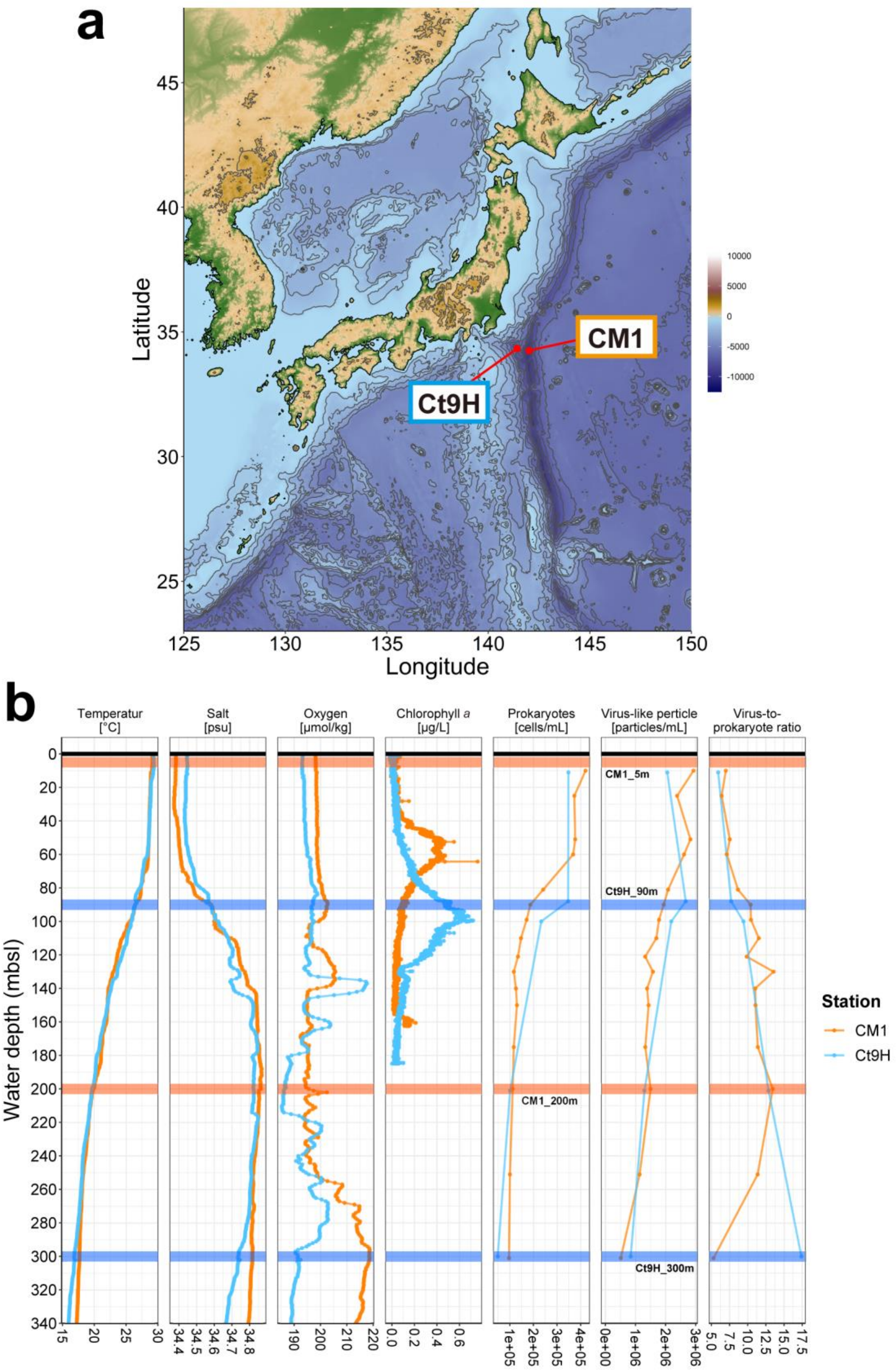


**Figure S1.** Hydrographic characteristics of the sampling sites. **a** Map of the Japanese archipelago. The red points indicate the sampling sites. **b** Vertical profiles of chemical concentrations and abundances of prokaryotic cells and virus-like particles. Profiles of water temperature, salinity concentration, dissolved oxygen concentration, and chlorophyll *a* concentration were measured *in situ*. Orange and blue represent CM1 and Ct9H sampling sites, respectively. The horizontal lines represent depths where the seawater samplings were conducted; 5 and 200 mbsl at CM1 site, and 90 and 300 mbsl at Ct9H site.

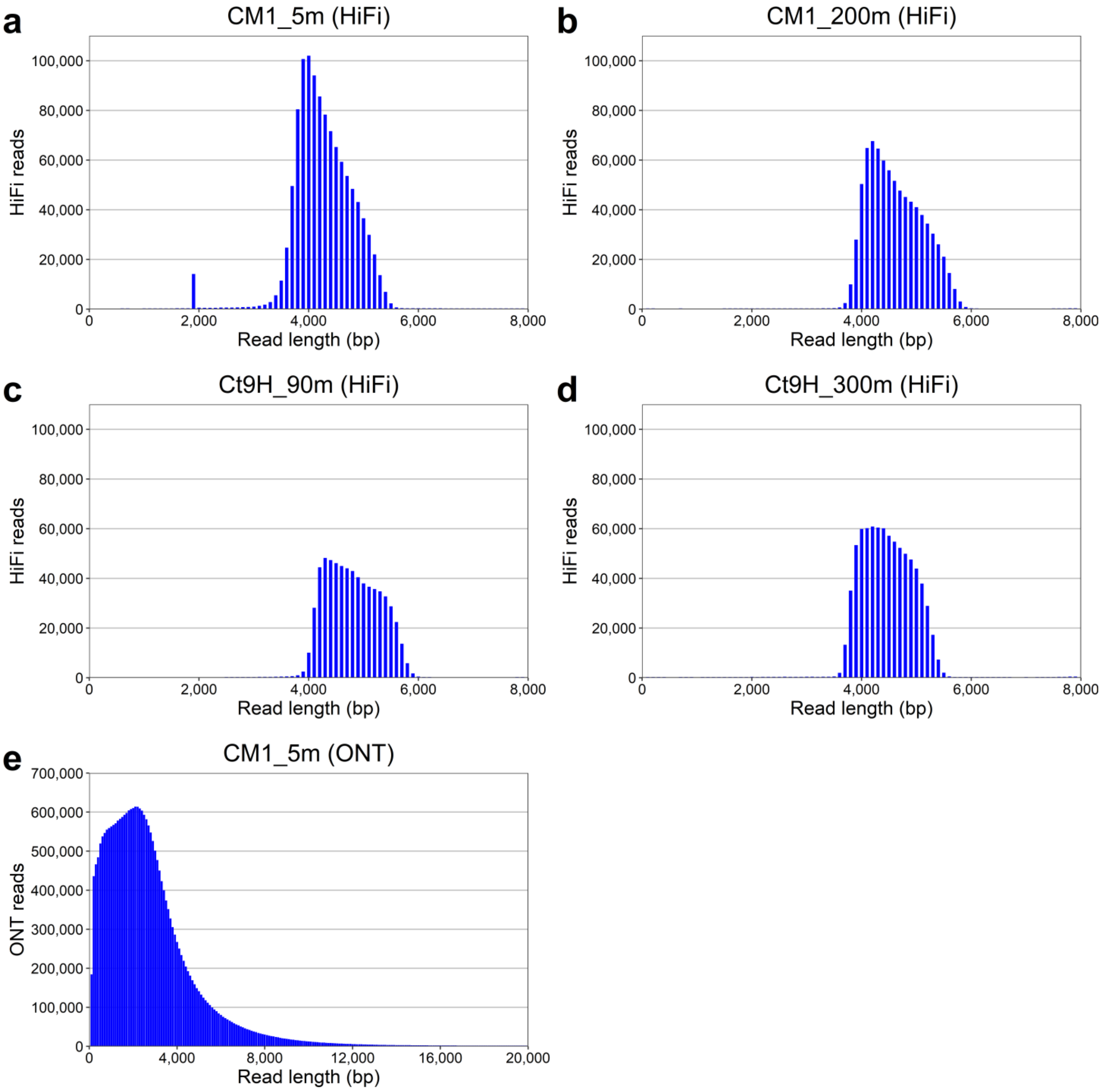


**Figure S2.** Frequency distribution of the length of sequencing reads. Read lengths were binned in 100-bp increments. **a**–**d** HiFi reads from PacBio Sequel. The SMRT libraries were size-selected at a 5-kb length. **a** We noted a small peak at 2 kb in CM1_5m, which likely reflected SMRT bell templates that unexpectedly collapsed either of the terminal hairpin adapters, although this may not significantly affect further sequencing reads and modification analyses. **e** ONT reads from Oxford Nanopore GridION. Reads ranging from 0–20 kb in length are presented. ONT libraries with short DNA fragment sizes were filtered out through the AMPure purification process in advance of the sequencing.

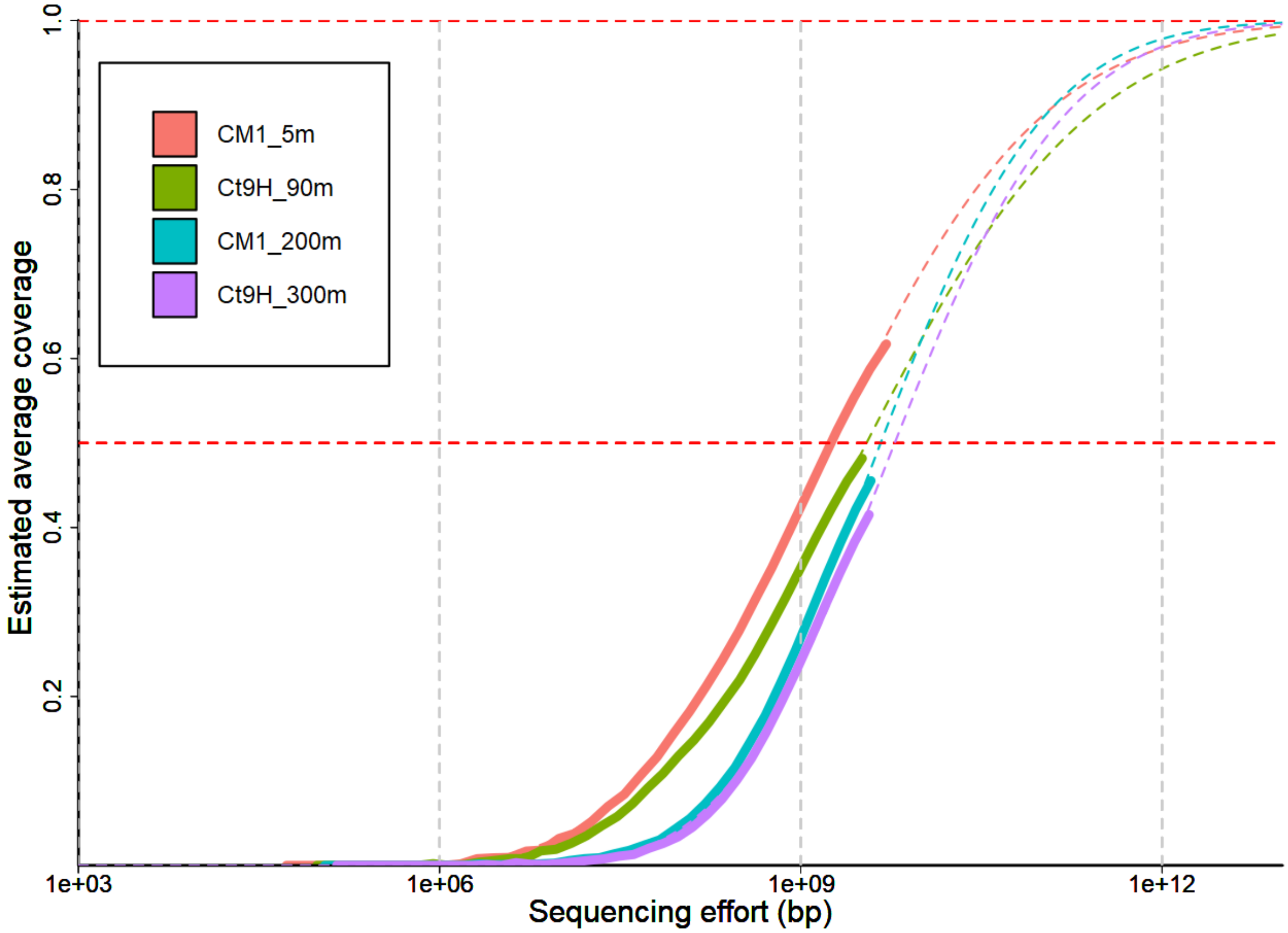


**Figure S3.** Estimated metagenomic coverage of HiFi reads. The dashed curves indicate the fitted models of the Nonpareil curves. The red horizontall dashed lines indicate 50% and 100% coverage.

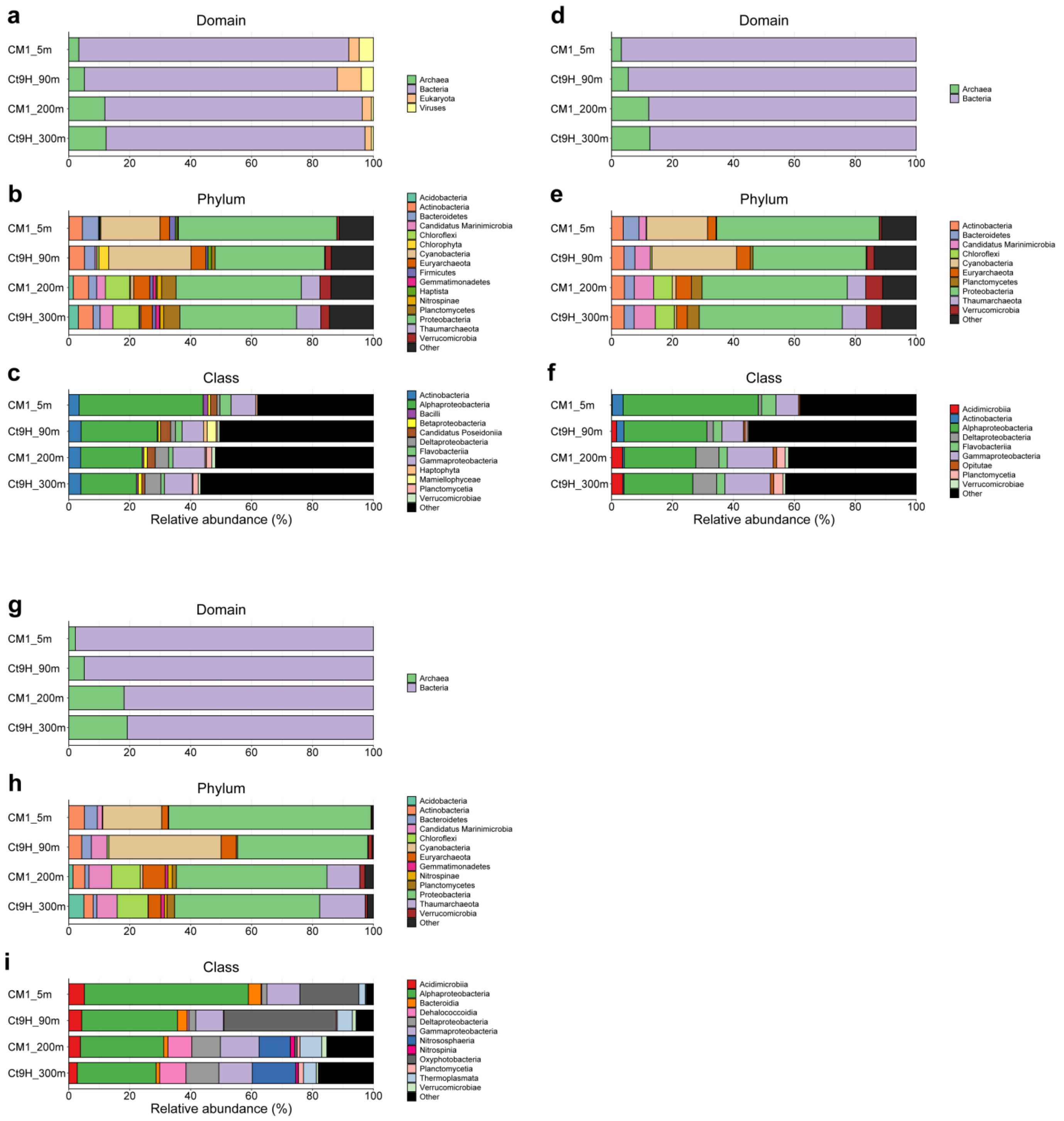


**Figure S4.** Taxonomic profiles of HiFi reads. Estimated relative abundances obtained using Kaiju with the NCBI nr database at the **a** domain, **b** phylum, and **c** class levels, those with the GORG database at the **d** domain, **e** phylum, and **f** class levels, and those from BLASTN analysis of full-length 16S rRNA gene sequences with the SILVA database at the **g** domain, **h** phylum, and **i** class levels are presented. The eukaryotic and viral reads are ignored in **d**-**i** based on the databases used. Sequences assigned to members with <1% abundance or taxonomically unclassified are grouped as ‘Others.’

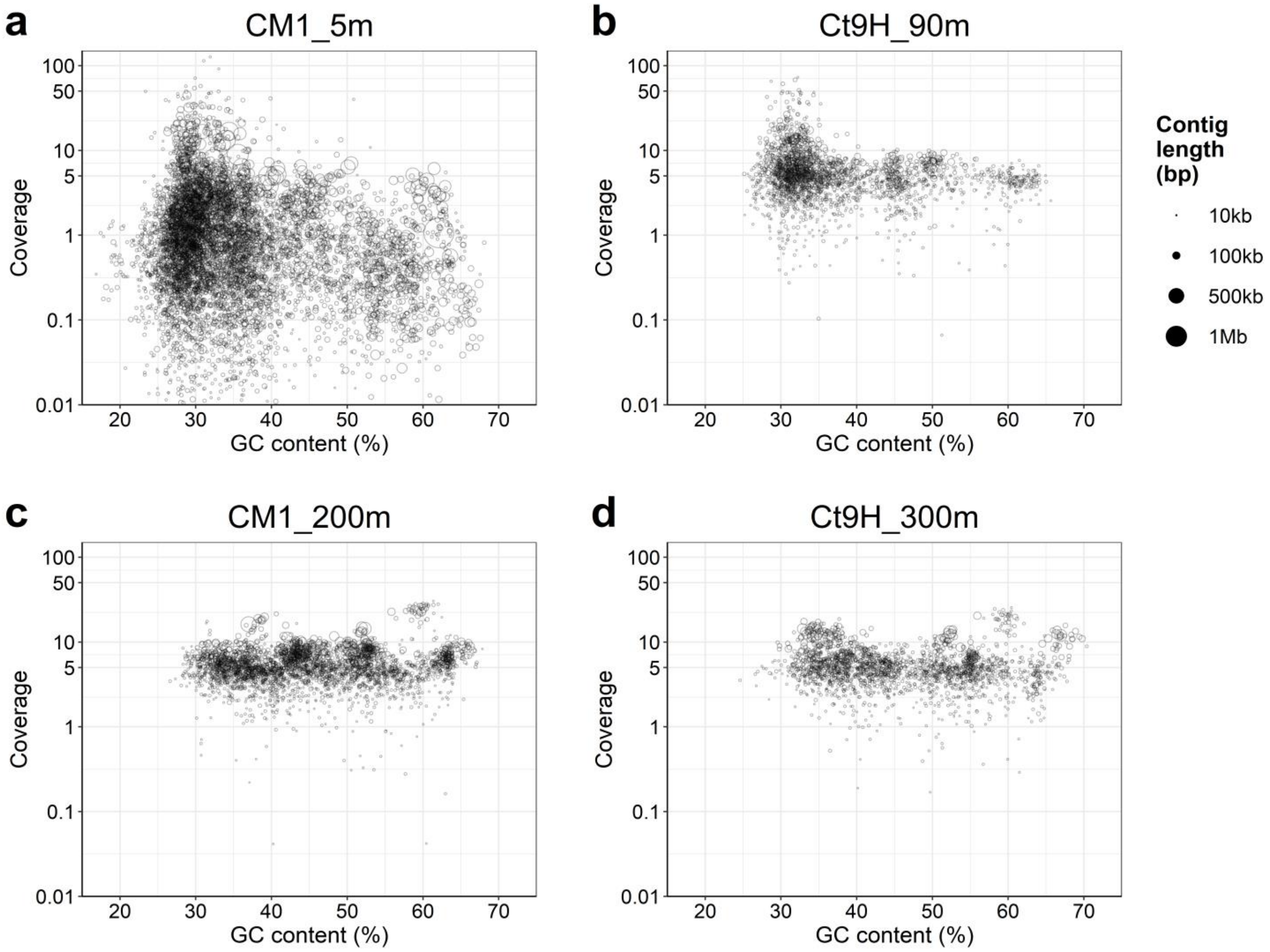


**Figure S5.** Assembled contigs. **a** CM1_5m assembled from ONT reads after polishing. Contigs with > 0.01× coverage against HiFi reads are presented. **b** Ct9H_90m assembled from HiFi reads. **c** CM1_200m assembled from HiFi reads. **d** Ct9H_300m sample assembled from HiFi reads. Each circle represents a contig, where the size represents its total sequence length. The x- and y-axes represent GC content and genome coverage of HiFi reads, respectively.

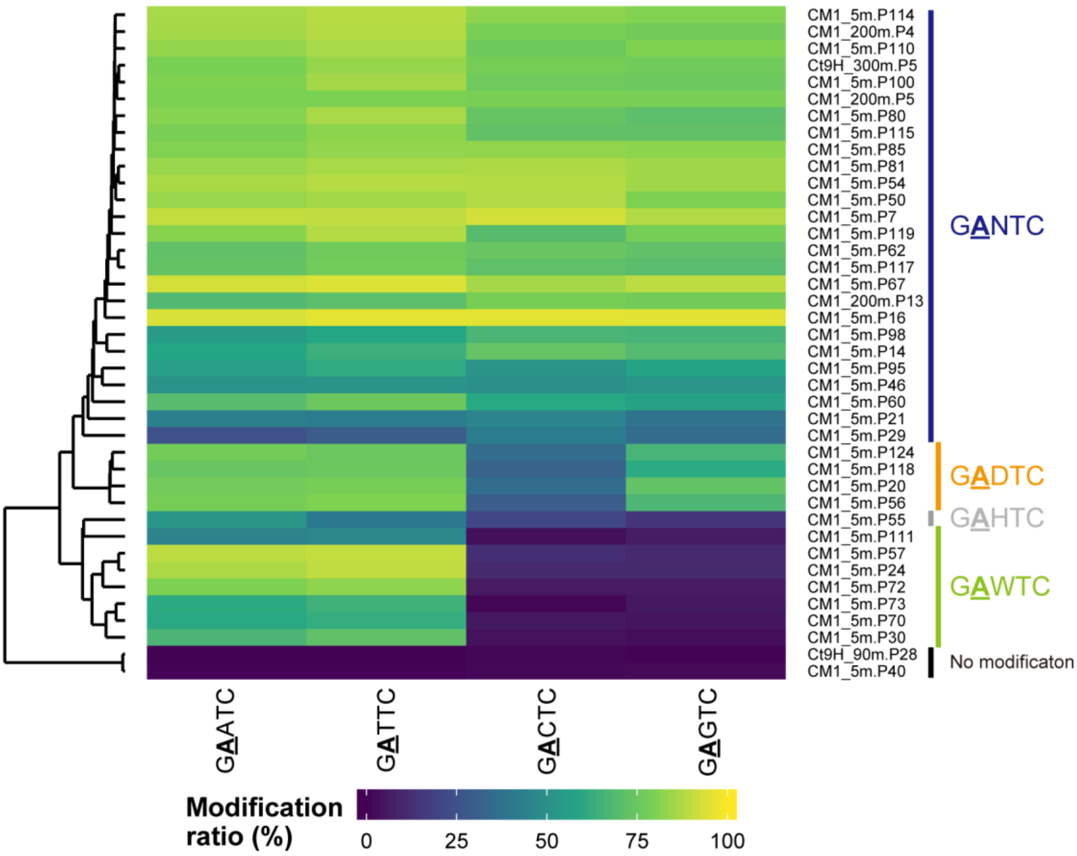


**Figure S6.** Methylomes of Alphaproteobacteria P-MAGs. The components of G**A**NTC are presented. The left dendrogram represents a hierarchical clustering of the modification ratios on each P-MAG. The distance matrix was calculated based on the Euclidean distance, and the clusters were calculated using the single-linkage method. Methylated motifs detected in each P-MAG using the metaepigenomic analysis are indicated on the right side. We noted that the G**A**HTC from CM1_5m.P55 was supported by low subread coverage (36×) and may be a misdetection.

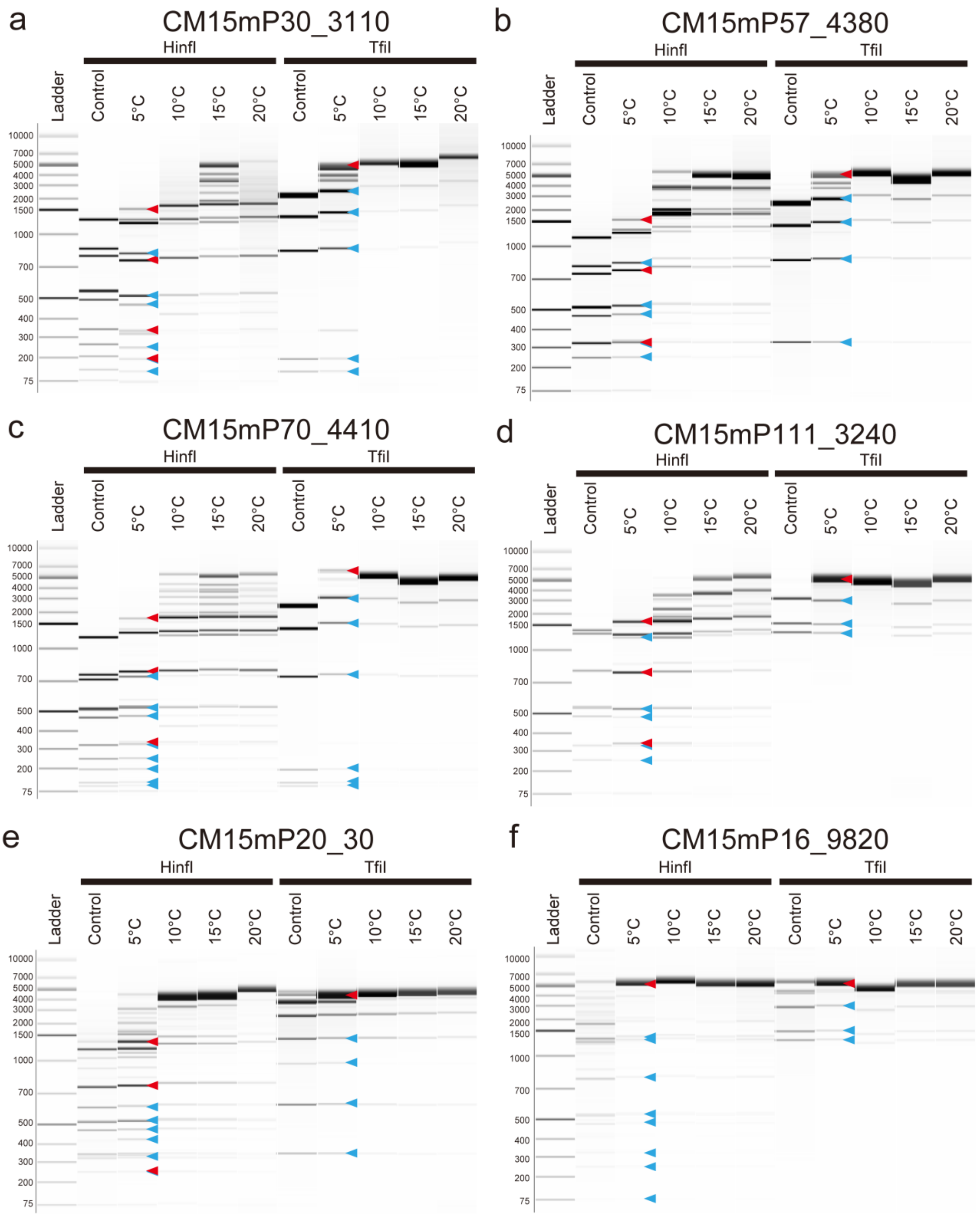


**Figure S7.** REase digestion assays of MTases that were detected in Alphaproteobacteria P-MAGs. HinfI (GANCT) and TfiI (GAWTC) REases were employed for the assays, where the plasmids contained 10–14 GANTC and 3–6 GAWTC target sites. All plasmid DNAs were linearized using NdeI. Based on the metaepigenomic analysis, **a-d** G**A**WTC, **e** G**A**DTC, and **f** G**A**NTC were hypothesized motifs of each MTase. Band sizes that theoretically appeared or were depleted when the hypothesized methylation occurred are indicated by red and blue triangle marks, respectively. We noted that several unexpected bands appeared in the upper side of the control samples of each HinfI and TfiI cleavage assays in **e** and **f**, possibly due to the leakage expressions from the constructs.

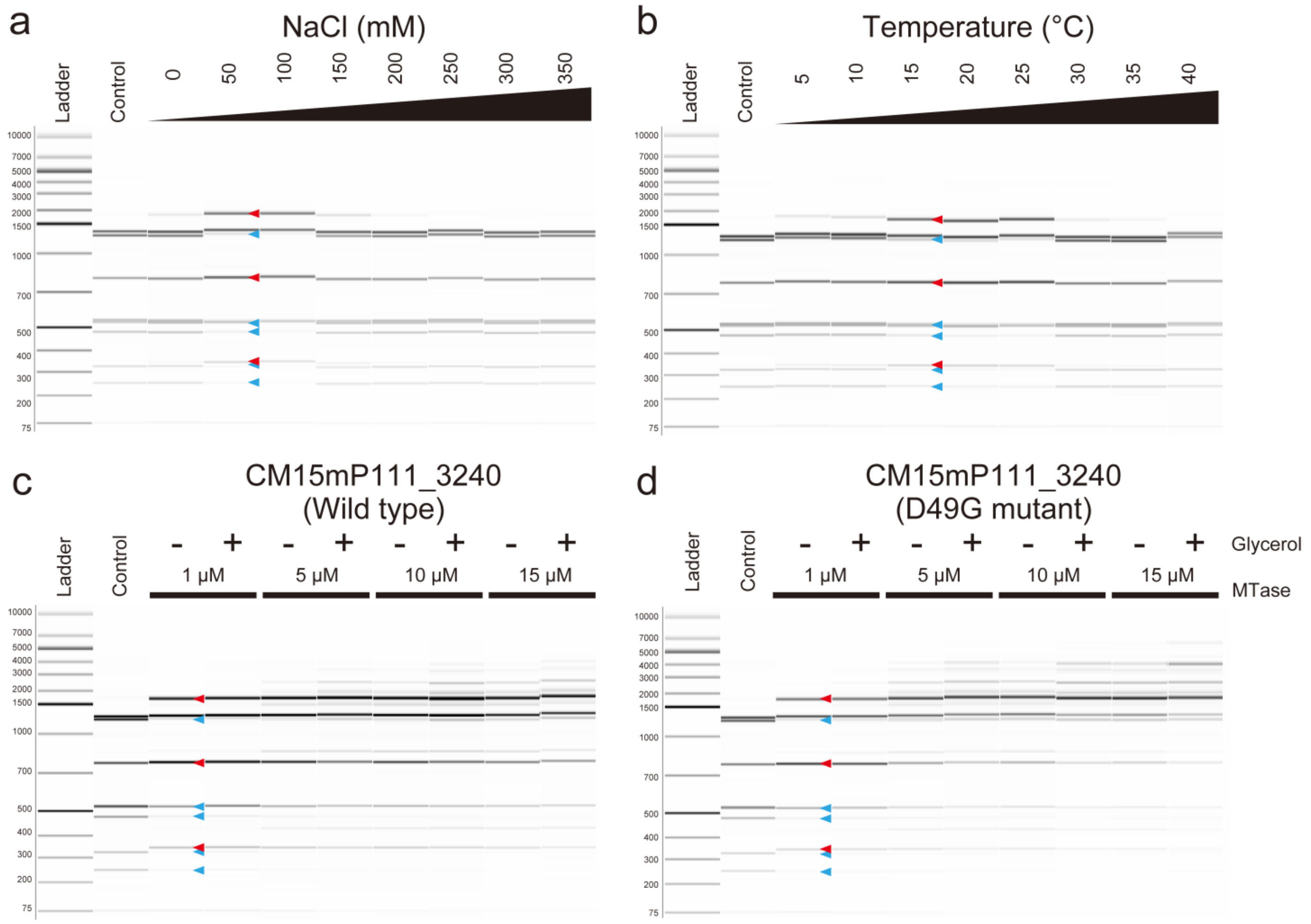


**Figure S8.** *In vitro* REase digestion assays of purified CM15mP111_3240 MTase and its mutant. Sensitivity assays of **a** salt and **b** temperature of the original MTase and assays of high enzymatic and glycerol concentrations in the **c** MTase and **d** D49G mutant are presented. Glycerol concentrations were changed to 0% (-) and 10% (+) (v/v). HinfI (GANCT specificity) was employed for the assays, where the substrate DNAs contained 10 GANTC target sites. Unmethylated substrate DNAs were produced using PCR. Band sizes that were expected to appear (red) and disappear (blue) when GA**W**TC methylation occurs are indicated with triangle marks. Substrate DNAs without MTase reaction were used for the control.

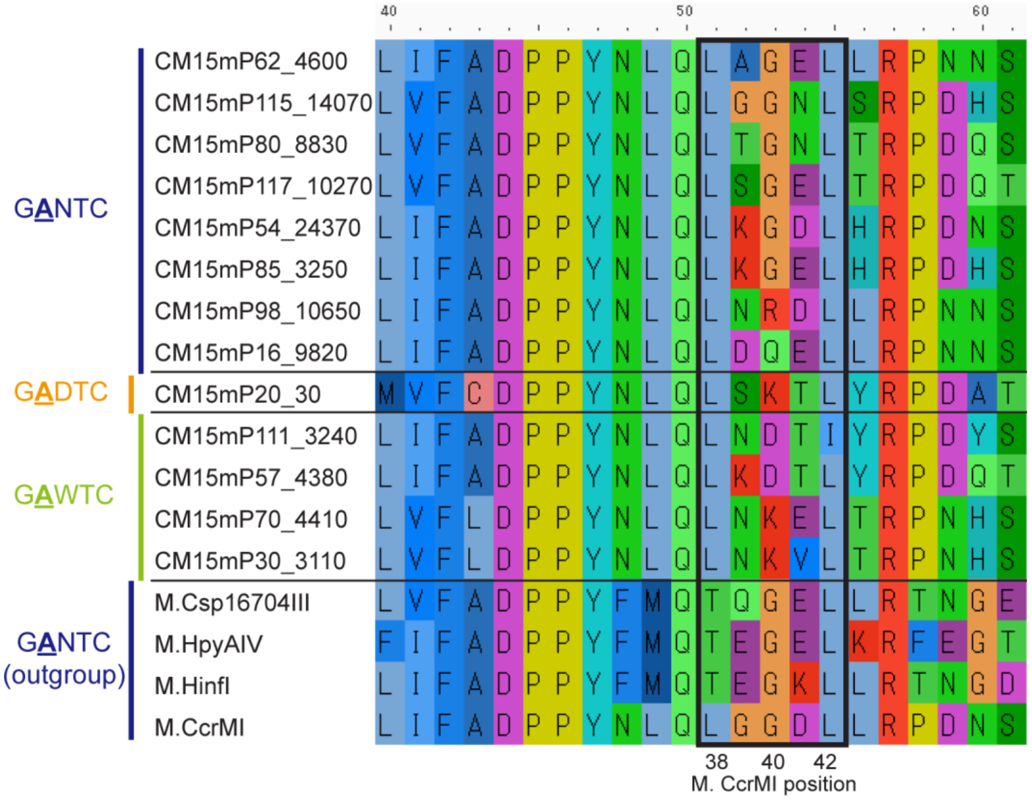


**Figure S9.** A multiple sequence alignment of M.CcrMI homologs. MTases that were predicted or experimentally confirmed to recognize either G**A**NTC/G**A**DTC/G**A**WTC were used in this analysis. The black rectangle indicates the region where they interact with the third position of the motif sequences (i.e., 'N' and 'W') in the M.CcrMI gene.

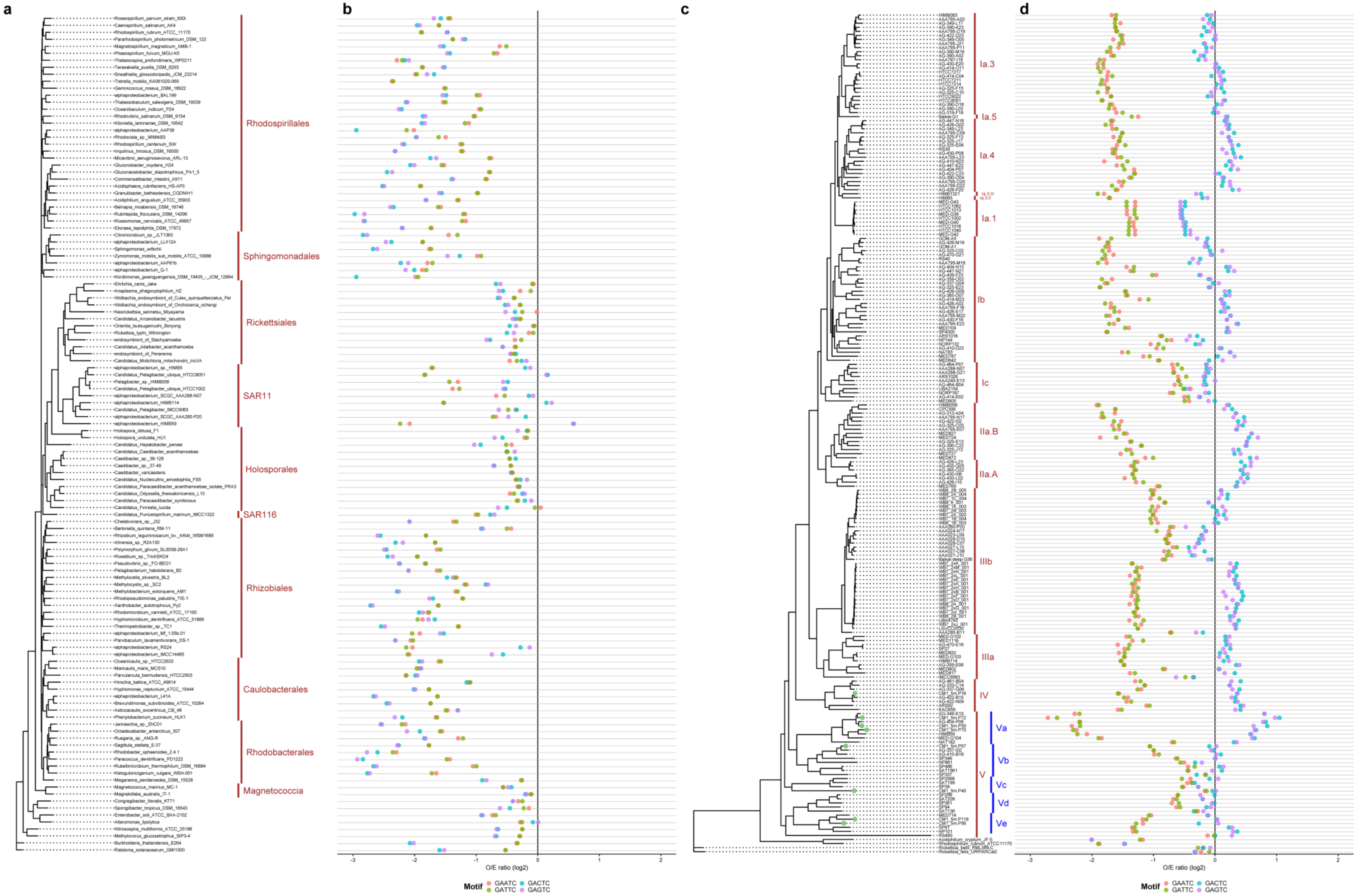
a
b
c
d
Rhodospirillales
Sphingomonadales
Rickettsiales
SAR11
Holosporales
SAR116
Rhizobiales
Caulobacterales
Rhodobacterales
Magnetococcia
Ia.3
Ia.5
Ia.4
Ia.1
Ib
Ic
IIa.B
IIa.A
IIIb
IIIa
IV
V
Va
Vb
Vc
Vd
Ve
O/E ratio (log2)
Motif
GAATC
GATTC
GACTC
GAGTC

**Figure S10.** Phylogenetic diversity of the Observed/Expressed (O/E) ratio of GANTC sequence. Pairs of a phylogenetic tree and O/E ratios of **ab** Alphaproteobacteria and **cd** SAR11 are presented. **a** Four Betaproteobacteria and four Gammaproteobacterial genomes were used as outgroups. Taxonomic groups at the order level are indicated in brown. We noted Magnetococcia (comprised of *Magnetococcus* and *Magnetofaba)* is currently classified as Alphaproteobacteria but was recently proposed to establish a novel Magnetococcia class. **c** The P-MAGs obtained in this study are colored in light green. One *Acidiphillum*, one *Rhodospirillum*, and two *Rickettsia* genomes were used as outgroups. Subclades are indicated in brown, and the inferred minor subgroups in subclade V referred to in this study are indicated with blue bars and texts. All the genomes retrieved from the NCBI repository are summarized in Supplementary Data S5.

### Supplementary tables

**Table S1.** Descriptions of sampling sites

| Sample | Cruise ID | Sampling method | Collection date | Area | Description | Sation | Location | Environmental feature | Elevation (mbsl) | Sample amount (L) |
|---|---|---|---|---|---|---|---|---|---|---|
| CM1_5m | KM19-07 | Intake pipe | 2019-09-02 | Izu-Ogasawara Trench | CM1 | 34N11 | 34.2607 N 142.0203 E | Epipelagic zone | 5 | 200 |
| CM1_200m | KM19-07 | Niskin sampler | 2019-09-02 | Izu-Ogasawara Trench | CM1 | 34N11 | 34.2607 N 142.0203 E | Mesopelagic zone | 200 | 180 |
| Ct9H_90m | KM19-07 | Niskin sampler | 2019-09-06 | Izu-Ogasawara Trench | Ct9H | 34N12 | 34.3317 N 141.4143 E | Epipelagic zone | 90 | 50 |
| Ct9H_300m | KM19-07 | Niskin sampler | 2019-09-06 | Izu-Ogasawara Trench | Ct9H | 34N12 | 34.3317 N 141.4143 E | Mesopelagic zone | 300 | 320 |

**Table S2.** Data on metagenomic shotgun sequencing read and HiFi read analyses

| | Sample | CM1_5m | Ct9H_90m | CM1_200m | Ct9H_300m |
|---|---|---|---|---|---|
| PacBio Sequel | Subread reads | 20,889,048 | 15,827,638 | 18,959,396 | 19,589,319 |
| | ---Read length (bp) | 5,001 ± 2,522 | 4,716 ± 1,963 | 4,519 ± 1,749 | 4,381 ± 1,708 |
| | ---Total base (bp) | 104,456,419,820 | 74,635,717,619 | 85,686,708,868 | 85,823,726,072 |
| | HiFi reads | 1,106,089 | 657,109 | 812,032 | 809,717 |
| | ---Read length (bp) | 4,311 ± 540 | 4,926 ± 771 | 4,707 ± 664 | 4,559 ± 630 |
| | ---Total base (bp) | 4,768,807,324 | 3,237,145,123 | 3,821,978,925 | 3,691,721,393 |
| | ---16S rRNA | 1,773 | 1,296 | 1,543 | 1,408 |
| | ---CDSs | 7,514,379 | 5,109,850 | 5,585,360 | 5,369,598 |
| | --- ---MTase | 27,676 | 8,067 | 12,846 | 3,324 |
| Oxford Nanopore GridION | Sequenced reads | 24,562,005 | - | - | - |
| | ---Read length (bp) | 2,734 ± 2,013 | - | - | - |
| | ---Total base (bp) | 67,142,060,909 | - | - | - |
| Illumina MiSeq | Paired-end reads | 9,433,870 | 8,637,416 | 10,330,676 | 9,015,972 |
| | ---Read length (bp) | 288 ± 31 | 286 ± 34 | 283 ± 34 | 284 ± 34 |
| | ---Total base (bp) | 2,720,391,360 | 2,472,592,381 | 2,919,215,542 | 2,558,126,139 |

**Table S3.** Data on metagenomic assembly and genome binning

| Sample | Contigs | Total length (bp) | N50 (bp) | Longest length (bp) | Average length (bp) | P-MAGs | Mapped HiFi reads on P-MAGs (%) | V-MAGs | Mapped HiFi reads on V-MAGs (%) |
|---|---|---|---|---|---|---|---|---|---|
| CM1_5m | 29,391 | 524,431,517 | 29,344 | 2,239,202 | 17,843 | 130 | 31.5 | 159 | 4.72 |
| Ct9H_90m | 7,829 | 82,604,737 | 11,600 | 307,709 | 10,551 | 30 | 9.4 | 3 | 0.19 |
| CM1_200m | 10,824 | 123,124,031 | 12,650 | 545,541 | 11,375 | 41 | 24.7 | 1 | 0.01 |
| Ct9H_300m | 8,145 | 82,769,163 | 10,530 | 362,843 | 10,162 | 32 | 19.9 | 0 | NA |

**Table S4.** Methylated motifs detected in the chromosomal DNA of *E. coli* transformed with artificially-synthesized MTase genes

| Genome ID | Transformed MTase gene ID | Detected methylated motif | Modification Type | Number of methylated sites | Number of motif sequences | Methylation ratio (%) | Mean modification QV | Mean subread coverage | Description |
|---|---|---|---|---|---|---|---|---|---|
| Ct9H_300m.P26 | Ct9H300mP26_1870 | BAAA**A** | m6A | 45818 | 47616 | 96.2% | 453.2 | 672.4 | |
| Ct9H_90m.P5 | Ct9H90mP5_10800 | BAAA**A** | m6A | 4375 | 47616 | 9.2% | 550.6 | 666.5 | |
| CM1_200m.P2 | CM1200mP2_32760 | CAA**A**T | m6A | 9630 | 9958 | 96.7% | 564.7 | 668.7 | |
| CM1_5m.P129 | CM15mP129_7780 | ACAA**A** | m6A | 10950 | 11372 | 96.3% | 459.4 | 675.0 | |
| CM1_200m.P10 | CM1200mP10_13750 | **C**TCC | m4C | 10992 | 19137 | 57.4% | 226.6 | 649.6 | |
| CM1_5m.P20 | CM15mP20_30 | G**A**DTC | m6A | 15451 | 17919 | 86.2% | 416.8 | 679.0 | Incubated at 5°C |
| CM1_5m.P20 | CM15mP20_30 | G**A**NTC | m6A | 9839 | 21484 | 45.8% | 535.8 | 681.1 | Incubated at 15°C |

R= A/G, Y= C/T, M= A/C, K= G/T, S= C/G, W= A/T, H= A/C/T, B= C/G/T, V= A/C/G, D= A/G/T, N= A/C/G/T
